# The actin cytoskeleton plays multiple roles in structural color formation in butterfly wing scales

**DOI:** 10.1101/2023.06.01.542791

**Authors:** Victoria J. Lloyd, Stephanie L. Burg, Jana Harizanova, Olivia Hill, Juan Enciso-Romero, Rory L. Cooper, Silja Flenner, Elena Longo, Imke Greving, Nicola J. Nadeau, Andrew J. Parnell

## Abstract

Vivid structural colors in butterflies are caused by photonic nanostructures scattering light. Structural colors evolved for numerous biological signaling functions and have technological applications. Optically, such structures are well understood, however their development *in vivo* remains obscure. We show that actin is intimately involved in structural color formation in the butterfly *Heliconius sara*. Using comparisons between iridescent (structurally colored) and non-iridescent scales in adult and developing *H. sara*, we show that iridescent scales have more densely packed actin bundles leading to an increased density of reflective ridges. Super-resolution microscopy revealed that actin is repeatedly re-arranged in later development, when optical nanostructures are forming. Furthermore, actin perturbation experiments at these later developmental stages resulted in near total loss of structural color. Overall, this shows that actin plays vital templating roles during structural color formation in butterfly scales, with mechanisms potentially universal across lepidoptera.

**Teaser:** The actin cytoskeleton is essential for templating the optical nanostructures responsible for structural color production in butterfly scales.

Actin templates the reflective ridges on butterfly scales and is directly involved in forming the color-producing nanostructures within these

## Introduction

Structural color produced by the interaction of light with nanostructures enables a diverse and tremendously vivid array of colors (*1*). They are particularly important in low light environments, for example in the forest understory, as they achieve superior visual signal propagation over pigmentary color (*2*). Despite the importance of biological photonic nanostructures from an evolutionary perspective and as designs for advanced optical materials (*3–5*), their structural formation remains poorly understood.

Photonic nanostructures within the wing scales are responsible for the structural color seen in butterflies and moths (*6*) these include; photonic crystals (*7, 8*), multilayer (Bragg) reflectors (*9*) and thin-films (*10, 11*). Each wing scale develops from a single cell, forming a chitinous envelope with an undifferentiated lower layer and a complex structured upper layer covered in longitudinal parallel ridges (*12–14*). In numerous structurally-colored butterfly species, these ridges are composed of multiple layers (lamellae), giving rise to constructive interference (*15–18*). Ghiradella (*16*) postulated that developing ridges buckle due to intracellular stress, and this is responsible for layered lamellae formation.

Studying the actin cytoskeleton during scale formation may improve our understanding of how layered lamellae form, as for many cell types it plays an important role in controlling cell shape (*19*). The scale ridges (on which the layered lamellae form) are the result of chitin deposition between parallel actin bundles (*13, 20, 21*). The actin bundles are temporary, and stabilized through polymerization and cross-linking of F-actin within developing scale cells (*22–24*).

The actin cytoskeleton in *Drosophila* bristles, a homologous structure to butterfly scales, has been extensively studied (*25*). Genetic knockouts of actin organization proteins have shown the actin cytoskeleton is important in controlling the number and shape of ridges in bristles, as well as localization of chitin synthetase enzymes, required to deposit the ridges (*23, 26*–*28*). In butterfly scales, the actin bundles may not just be limited to guiding ridge positioning but could be crucial in sculpting finer-scale aspects of scale morphology, including the photonic nanostructures.

*H. sara,* along with several closely related species in the same clade, are fairly unusual in the *Heliconius* genus in displaying iridescent blue wing coloration (*29–31*)(Fig 1A-1B). *H. sara* has both structurally colored blue iridescent and non-structurally colored black scales (Fig 1A-1C), facilitating direct comparisons of these scale types throughout their development. The structural color of *H. sara* is generated through layered lamellae on the parallel scale ridges (Fig 1F-1G) (*29, 30*).

**Fig 1:**
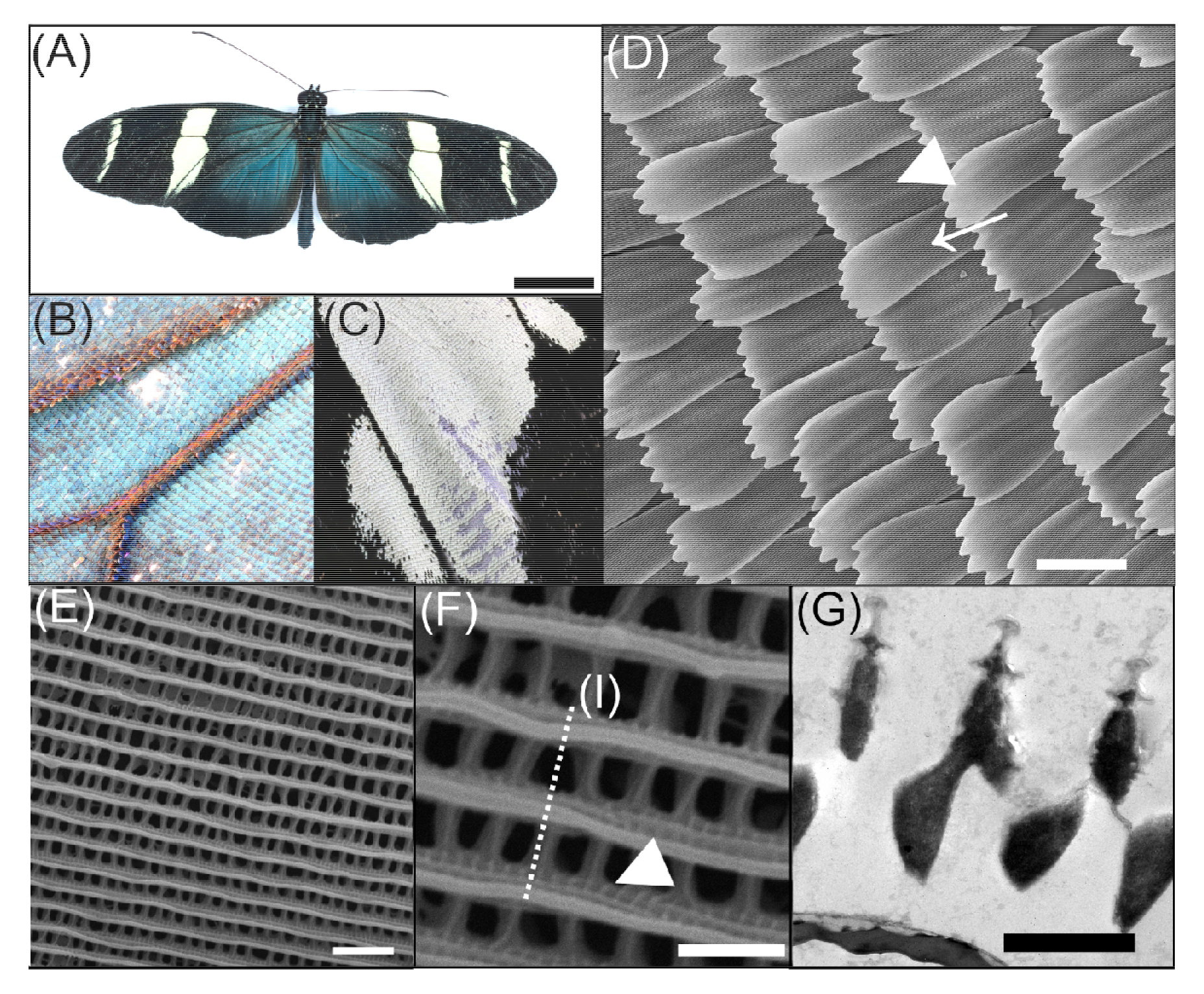
The neotropical butterfly *Heliconius sara*. **(A)** Dorsal view of a *Heliconius sara* individual. (B) Region of blue, iridescent wing scales on the proximal forewing. (C) Region of black and white, non-iridescent wing scales on the distal forewing. (D) SEM image of the overlapping scales on the dorsal wing surface. Cover scales (arrow) sit directly on top of the basal ground scales (arrowhead). (E) Dorsal view of an iridescent wing scale surface, with many periodically ordered longitudinal ridges running parallel to scale length. (F) High-magnification view of an iridescent wing scale showing ridge ultrastructure; with open windows into the scale lumen separated by crossribs. Microribs (arrowhead) pattern the sides of the ridges and are perpendicular to ridge direction (dashed line). (G) Transmission electron microscope (TEM) cross-section through the scale ridges. The layers on the ridges form a multilayer photonic nanostructure. Scale bars lengths: (A) 10 mm, (D) 50 µm, (E) 2 µm, (F, G) 1 µm.

Here, we examine F-actin organization during wing scale development in the butterfly *H. sara*, focusing on the formation of the nanostructures responsible for iridescence. Using scanning electron microscopy (SEM) and fluorescence microscopy we investigate whether patterning of F-actin differs between iridescent and non-iridescent wing scales. We use lifetime separation stimulated emission depletion (TauSTED) super-resolution microscopy (*32*) to gain insight into actin remodeling during scale development. We then chemically perturb actin dynamics to elucidate whether the actin cytoskeleton plays a direct role in the formation of optical nanostructures in *H. sara*.

## Results

To compare the adult morphology of iridescent and non-iridescent scales on the dorsal forewing of *H. sara* we used SEM and X-ray tomography (Fig 2, S1, S2 Movie, S3 Movie, S4). We examined both upper cover scales and basal ground scales (Fig 2D). The general structure of iridescent and non-iridescent scales is almost identical (S2, S3), with both having a flat smooth lower layer (lamina) and a highly intricate upper layer (lamina). The parallel ridges on the upper lamina are joined together by crossribs, with the spaces between crossribs forming a regular series of windows into the interior scale lumen (Fig 2C).

**Fig 2:**
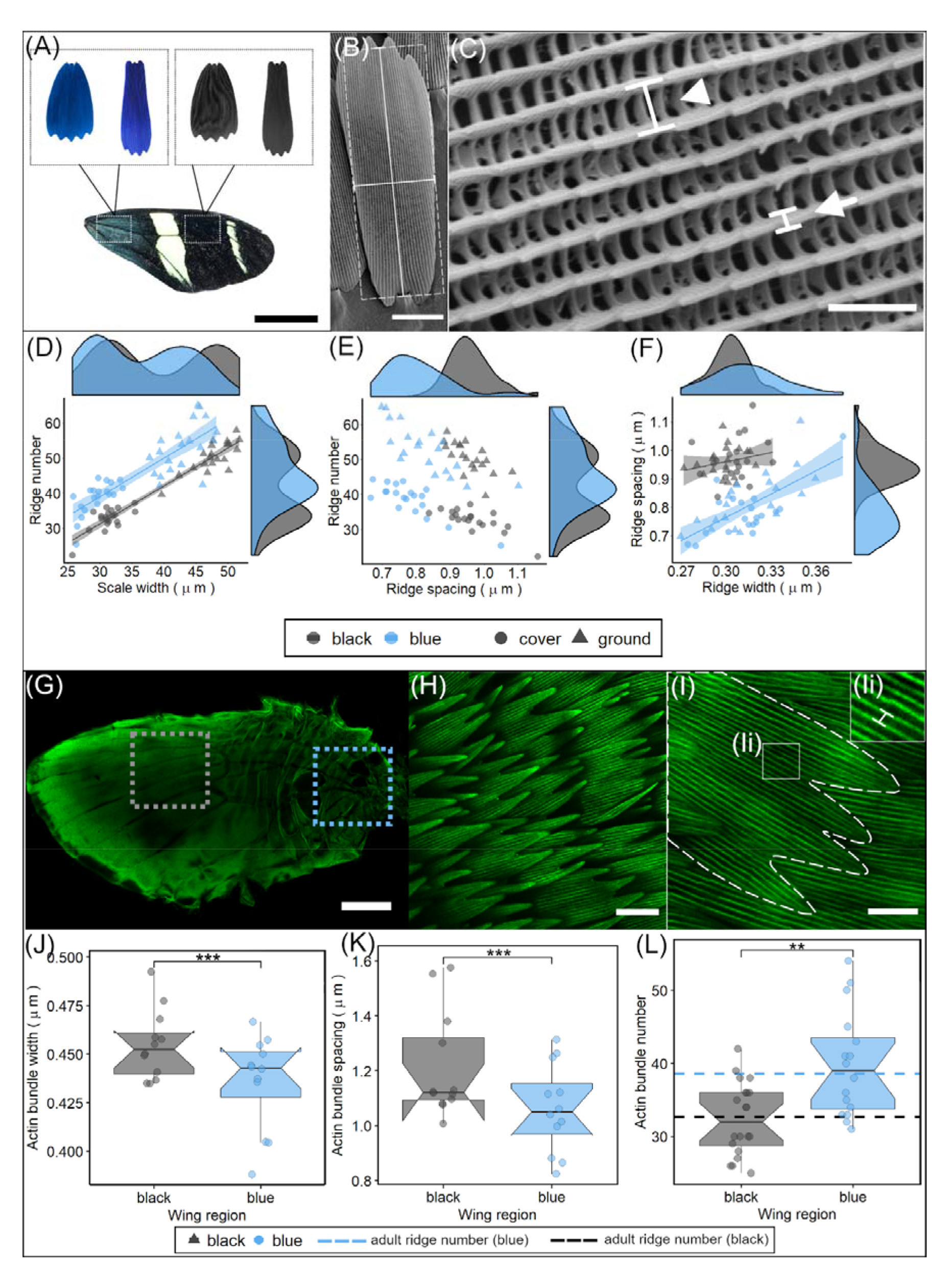
Morphological analyses of adult ridge organization and pupal actin patterning. A) Cover and ground scales (SEM images, shown in false color) were sampled from the proximal, iridescent (blue) wing region and the distal, non-iridescent (black) wing region. Representative SEMs showing measurements of (B) scale length (vertical solid line), width (horizontal solid line) and approximate area (dashed line); and (C) ridge spacing (arrowhead) and ridge width (arrow). Comparison of cover and ground scales in blue and black wing regions for (D) ridge number and scale width (µm) (E) number and ridge spacing (µm) (F) ridge spacing (µm) and ridge width (µm). Each point is the mean value grouped by individual, region and scale type. Shaded areas around regression lines indicate 95 % confidence intervals. Density plots on the axes give the distribution of each parameter for iridescent and non-iridescent scales separately (cover and ground combined). G) whole-mounted phalloidin-stained *H. sara* forewing, showing the iridescent region (blue box) and non-iridescent region (gray box). H) overlapping wing scales at 50%, with actin bundles visualized through phalloidin staining. I) Extraction of measurements of actin bundles from an individual developing scale. Ii) High-magnification zoom of the individual actin bundles showing the spacing between two adjacent bundles. J) Actin bundle width (μm) for 50% iridescent (blue) and non-iridescent (black) scales. K) Actin bundle spacing (μm) for 50% iridescent (blue) and non-iridescent (black) scales. L) Actin bundle number for iridescent (blue) and non-iridescent (black) scales, dashed lines indicate ridge number in adult cover scales. Points in (J, K) represent mean measurements for each individual grouped by region, points in (L) represent individual scales. Scale bar lengths: (A) = 10 mm, (B, H) = 20 µm, (C) = 2 µm, (G) = 1 mm, (I) = 10 μm;.

Correlation function analysis of the X-ray nano-tomography measured scales indicates a greater crossrib spacing in the black scale compared to the iridescent scale (iridescent 0.483μm; non-iridescent 0.607μm)(S1) (*33*). An expanded crossrib spacing in black scales likely allows more light to enter the scale and so be absorbed by melanin pigments (*34*).

Both cover and ground iridescent scales were smaller in size than non-iridescent scales (mean ± SE scale area, blue: cover 2700μm^2^±21, ground 3708μm^2^±35; black: cover 3044±25μm^2^, ground 4123μm^2^±30; likelihood ratio, χ^2^=208, d.f. = 1, p<0.001), which can be attributed to the decreased width of iridescent scales (mean ± SE scale width, blue: cover 29.5μm±0.22, ground 42.9μm±0.3, black: cover 31.3μm± 0.22, ground 47.8 μm±0.33; likelihood ratio, χ^2^=24, d.f. = 1, p<0.001; Fig 2D, S5B).

Having confirmed that the general structure of iridescent and non-iridescent scales are similar we next quantified differences in the finer scale elements, focusing first on the parallel ridges (Fig 2C, S4). The iridescent blue scales had significantly reduced ridge spacing compared to the non-iridescent black scales (mean ± SE ridge spacing, blue 0.804μm±0.007, black 0.962μm±0.004; likelihood ratio, χ^2^=446, d.f. = 2, p<0.001; Fig 2E-F). The reduced ridge spacing in blue scales was also confirmed via correlation function analysis of the tomography data (S1E,F) (*33*). This within-wing difference is consistent with prior work comparing between species and populations, which found iridescent *Heliconius* species have reduced ridge spacing compared to non-iridescent species (S6) (*29*). The decreased ridge spacing in iridescent scales can be attributed to an overall increase in ridge number, rather than a smaller scale width, with iridescent scales consistently having a greater ridge number for a given scale width (Fig 2D). There was also an effect of scale type (cover or ground) on ridge spacing (likelihood ratio, χ^2^=27, d.f. = 2, p<0.001). Iridescent cover scales had significantly reduced ridge spacing compared to iridescent ground scales (Tukey comparison, p < 0.001), but there was no difference in ridge spacing between cover and ground scales for non-iridescent scales (Tukey comparison, p=0.633).

Ridge width was slightly greater in iridescent scales compared to non-iridescent scales (mean ± SE ridge width, iridescent 0.315μm±0.002, non-iridescent 0.302μm±0.001; likelihood ratio, χ^2^=43, d.f. = 1, p<0.001; Fig 2F). There was no difference in ridge width between cover and ground scales (likelihood ratio, χ^2^=0.24, d.f. =1, p=0.622, Fig 2F). Interestingly, the distribution of ridge width in iridescent scales was much greater than that of non-iridescent scales (S5C).

Whilst the general morphology of adult *H. sara* iridescent and non-iridescent scales is similar, morphological differences in the scale ridges are observed, with adult iridescent scales displaying slightly thicker ridges and considerably reduced ridge spacing compared to non-iridescent scales. We were not able to quantify differences in the layering of lamellae within the ridges, which has previously been shown to be responsible for the iridescent color (*29, 30*), as this was beyond the resolution of the x-ray nano-tomography and would have required TEM sections of the ridges. However, the increased density and number of ridges likely contributes to the increased reflectance of the iridescent scales (*29, 35*).

### Development of H. sara scales

We characterized scale development from 25% to 62.5% of pupation, this encompasses scales emerging from the wing epithelium to formation of the final scale morphology (Fig 3). Overall, the development of iridescent and non-iridescent scales was very similar, and comparable to that reported for other butterfly species (*21*). At approximately 25%, nascent scales begin to emerge, as small actin-dense cytoplasmic projections from the wing epithelium (Fig 3A). Scale cell nuclei sit directly within the wing epithelium and are considerably larger than surrounding nuclei. Alpha-tubulin staining at 31% reveals the emerging scale buds are rapidly filling with cytoplasm (Fig 3B,C) and beginning to differentiate into cover and ground scales, with the larger ground scales containing more cytoplasm. In some cases, the tubulin appears organized into dense arrays, suggesting ordered microtubules are beginning to form (Fig 3C). By 37.5% the scales are essentially elongated sacs, containing thick longitudinal actin bundles (Fig 3D). Previous research has shown that actin bundles are required for scale elongation. These form through polymerization of actin into filaments (F-actin), followed by cross-linking of filaments together into thick bundles (*20, 21*). The actin bundles are most clearly discernible at the proximal portion of the scale where it buds from the epithelium through the developing socket (Fig 3F).

**Fig 3:**
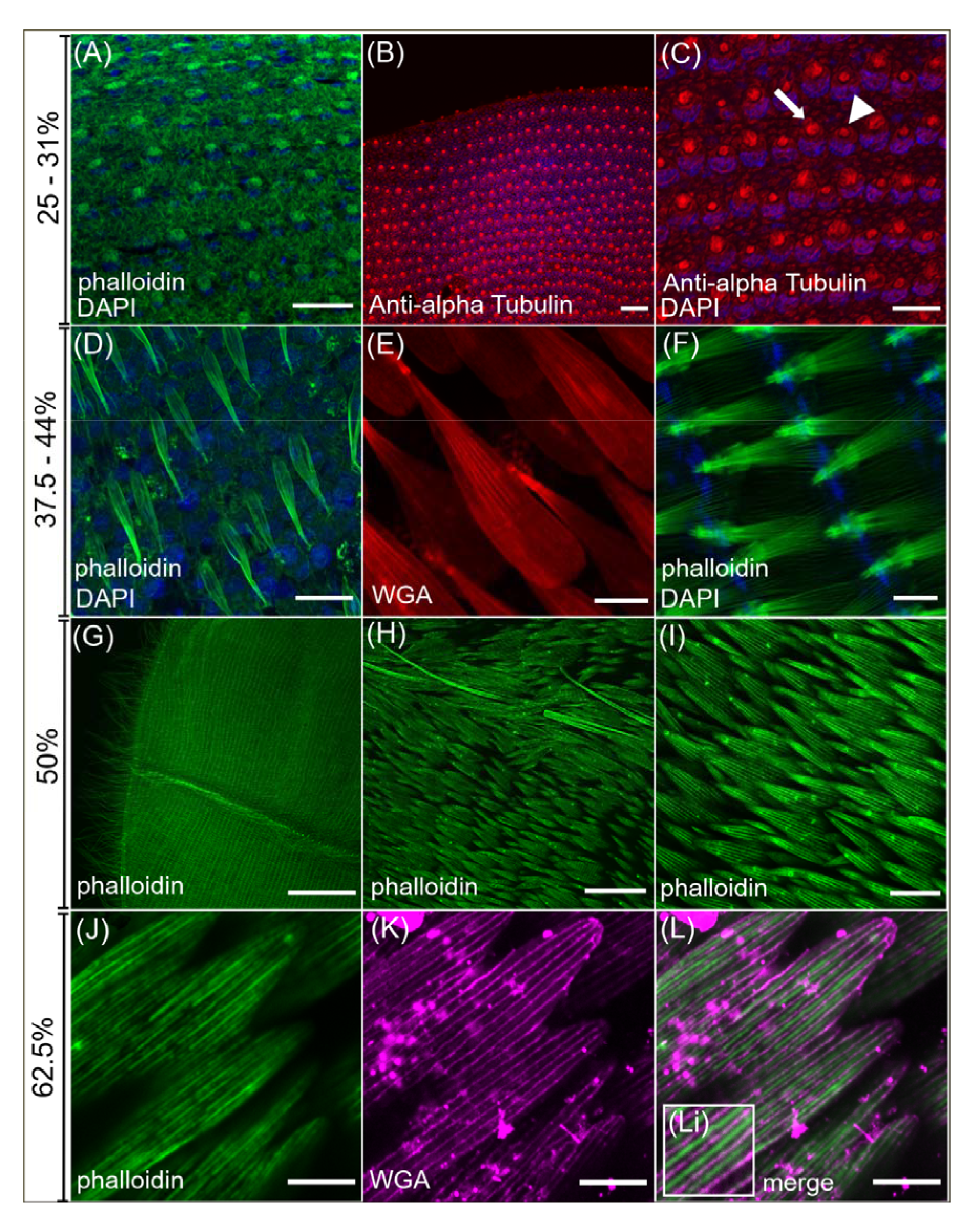
Confocal series of normal wing scale development in *H. sara.* Cell nuclei counterstained with DAPI (blue). A – C) Early wing scale development showing cytoplasmic projections from the wing epithelium at 25%. A) Phalloidin staining (green) of actin in the nascent scales. B, C) Anti-alpha Tubulin immunostaining (red) reveals differing amounts of cytoplasm in developing cover (arrowhead) and ground (arrow) scales and outlines of the socket cells. D – F) At 37.5–44% the scale cell is a sac filled with organised actin bundles (green) (D) and surrounded by a cellular membrane, highlighted by WGA staining (red) (E). Forming sockets are clearly visible (F) with the actin bundles passing directly through them. At 50%–56% (G - I) the scales resemble adult scales (Fig 1D). The distal forewing (G) shows hundreds of developing scales. H) overlapping wing scales adjacent to a wing vein with actin-rich hairs protruding from the vein (arrowhead). I) The actin (green) within the scales is highly organized at 50% and extends to the proximal portion of the scale fingers. J – L) final stages of scale development. J) gaps between the phalloidin stained actin bundles (green) highlights actin sub-bundling (K) WGA (magenta) now stains the chitin being deposited extracellularly (L) Merge of actin (green) and WGA (magenta) shows the chitin being deposited between the actin bundles (Li). Scale bar lengths: (A, B, E, I) 20μm; (C, D, F, J, K, L) 10μm; (G) 300μm; (H) 50μm.

At 50%-56% the scales become flattened and long finger-like projections form on the distal tip (Fig 3G-I). At this stage, the actin bundles are highly ordered in appearance and cover the entire proximal-distal portion of the scale (Fig 3I). At around 62.5%-69% chitin is deposited between the parallel actin bundles to form the cuticle ridges.

### F-actin patterning differs between developing iridescent and non-iridescent scales

We determined the optimal developmental stage to quantify actin organization as 50% of total pupal development. At this stage, actin bundles are highly regular and have reached the distal portion of the scale (Fig 2I, 3I) (*20, 21*). Additionally, chitin ridge deposition is beginning, suggesting that the actin bundles are correctly positioned for ridge formation to occur.

We quantified the spacing and thickness of actin bundles within developing scales using confocal microscopy of phalloidin-stained wings (Fig 2G-L). Iridescent scales had slightly thinner actin bundles compared to non-iridescent scales (mean ± SE bundle width, iridescent 0.438μm±0.004, non-iridescent 0.456μm±0.003; likelihood ratio, χ^2^=19, p<0.001; Fig 2J), although this may be influenced by slight differences in development stages observed between the proximal and distal forewing scales (*21*). The developing iridescent scales had reduced actin spacing compared to the non-iridescent, black scales (mean ± SE bundle spacing, iridescent 1.07μm±0.02, non-iridescent 1.22μm±0.03; likelihood ratio, χ^2^=40, p<0.001; Fig 2K). Furthermore, we found that iridescent scales had a greater number of actin bundles compared to non-iridescent scales (mean ± SE actin bundle number, iridescent 40±1.8, non-iridescent 32±1.1; likelihood ratio, χ^2^=11, p<0.001; Fig 2L)

This result is consistent with previous findings (*13, 21*), indicating a tight coupling between the spacing of actin bundles and spacing of chitin ridges, for both iridescent and non-iridescent scales. The mean number of actin bundles in iridescent and non-iridescent cover scales closely matched the mean number of ridges measured in adult cover scales of both types (Fig 2L). Our results show that the patterning of actin in developing *Heliconius* scale cells plays an important role in governing the density of adult scale ridges, which is an important morphological parameter controlling the iridescent properties.

### TauSTED super-resolution microscopy reveals detailed remodeling of the actin cytoskeleton

To investigate the ultrastructural remodeling of the actin cytoskeleton during the development of *H. sara* scales we used TauSTED microscopy (Fig 4,5, S8). At 44% of pupal development we observe both smaller peripheral actin bundles as well as larger internal actin bundles, described previously by Dinwiddie et al., (*21*) (Fig 4A-C). In scales with incipient finger formation, these were seen as vertices forming on the previously smooth distal edge, giving the tip of the scale a trapezoid-like shape. The finger origins coincide with the locations at which prominent internal actin bundles appear to attach to the distal membrane (Fig 4C, arrowheads; S7, animation). This hints at a possible role of these larger internal actin bundles in specifying spatial positioning of the fingers. Previous actin inhibition experiments performed by Dinwiddie et al., (*21*) resulted in scales lacking fingers, consistent with a role of actin bundles in specifying finger position and elongation.

**Fig 4:**
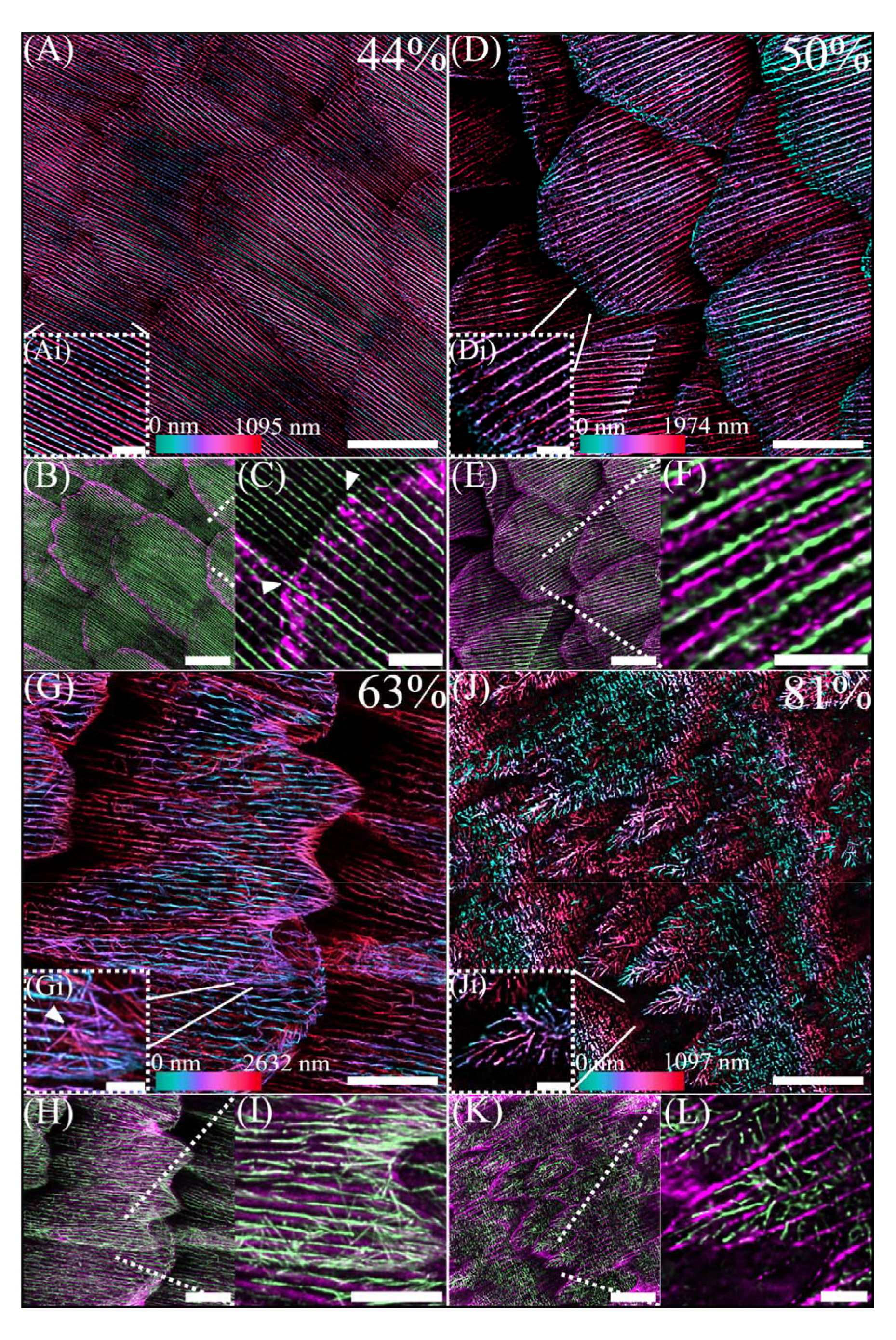
TauSTED super-resolution microscopy of the rearranging actin cytoskeleton during the development of *H. sara* scales. Depth colored images (A, D, G, J) show F-actin stained with phalloidin. Colored images below (B, C, E, F, H, I, K, L) represent a merge of actin (phalloidin, green) and chitin (CBD-TMR, magenta). A-C) At 44% of development small, numerous actin bundles are visible which extend to the distal edge of the cell. Color depth profiles (Ai) indicate smaller actin bundles are present on the dorsal surface of the cell (blue) whereas larger actin bundles are located more internally (red/magenta). Incipient cuticle formation begins at the periphery of the scale cells (B). Points of finger origination (arrows in C) correspond to locations of larger, internal actin bundles associating with the scale tip. D-F) At 50% the actin bundles are maximally spaced as the scale cell becomes increasingly flattened. Cuticle formation is evident across the scale (E), with cuticle ridges appearing in between actin filaments (F). G-I) At 63% the large continuous, parallel actin bundles are dissociating. A second network of branched F-actin is located more internally (blue in G) and is particularly evident along the scale edges and the fingers. Many of the individual filaments appear to radiate from single point of origination (arrow in Gi) and span across several microns before apparently attaching to the edge of the scale cell. (J-L) At 81% the actin network within the cell undergoes a final rearrangement. At the finger tips, highly branched actin projects from within the middle portion of the fingers towards the distal edges (Ji). Within the scale body no parallel bundles or branched filaments are visible, instead the actin has taken on ‘block’ like appearance. Ridge cuticle formation is complete and ultrastructures such as the crossribs are visible (K, L). Scale bars: (I) 5μm; (A, B, D, E, G, H, J, K) 10μm; (C, F, L, Ai, Di, Gi, Ji) 2μm.

**Fig 5:**
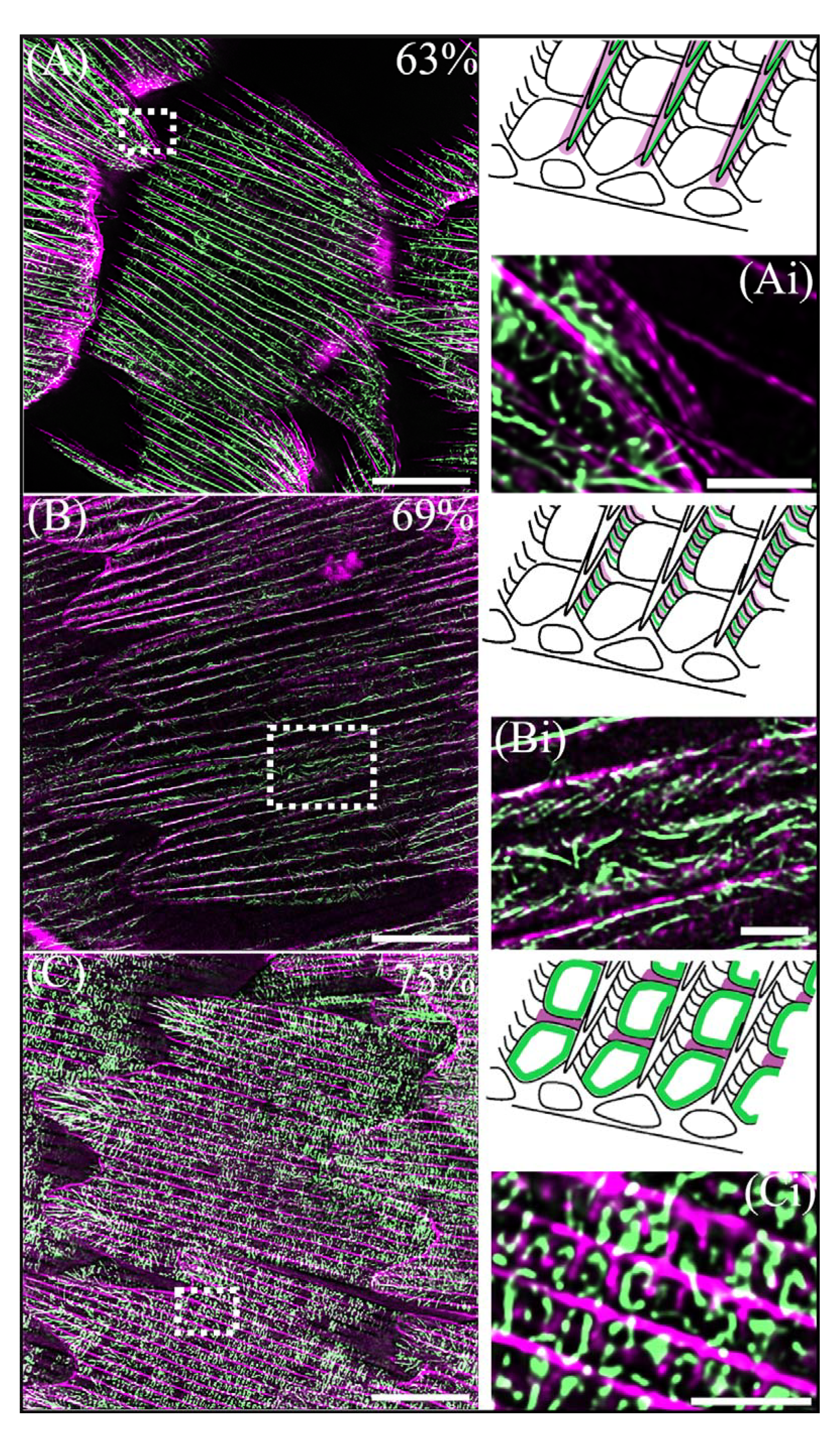
TauSTED super resolution imaging of actin filaments associated with various scale ultrastructures. Merge of actin (phalloidin, green) and chitin (CBD-TMR, magenta). Cartoon insets highlight the purported location of the actin filaments (green) and the associated cuticle structure (magenta). A) Black scale at 63% development, with Ai showing an enlarged view of a scale finger. The ridges at the edge of the scale finger are angled out of plane revealing the layering of a cuticle ridge and actin filaments below. B) Black scale at 69% development, with numerous individual actin filaments patterned along the side of a ridge (Bi), appearing similar to the microribs of adult scales. C) Iridescent scale at 75% development. The main scale body is filled with square ‘blocks’ of actin. The enlarged section (Ci) indicates the blocks of actin occur within the window regions in between the crossribs but do not fill these regions entirely. Scale bars: (A, B, C) 10 μm; (Ai, Bi, Ci) 2 μm.

At 50% the actin bundles are maximally spaced in agreement with our confocal microscopy observations (Fig 3). Z-stacks of the optical sections suggest re-structuring of the actin bundles, with the continuous uniform actin bundles, now displaying a more intricate ultrastructural arrangement (Fig 4D-F). In addition, some actin appears to be present between the large bundles (Fig 4F), reminiscent of the transient ‘actin snarls’ described in *Drosophila* bristle development (*23, 28, 36, 37*).

At 63% of development the large actin bundles are undergoing disassembly, with fracturing of the bundles into disjointed sub-bundles (Fig 4G-I). A previously undescribed second population of branched actin is now present and is particularly evident at the scale edges as well as the fingertips (Fig 4I). These branched actin filaments are smaller in diameter, located more internally (Fig 4Gi) and are orientated multi-directionally compared to the actin bundles. Along the scale edge, multiple filaments appear to radiate like spokes from single points further inside the scale that connect with the scale edge (Fig 4I).

At 75-81% the actin cytoskeleton undergoes a final, further reorganization with a highly branched network present in the fingers, radiating towards the distal fingertips (Fig 4J-L, 5C). In contrast, the main scale body is now devoid of any parallel actin bundles and is instead entirely filled with square ‘blocks’ of actin which run the length of the scale and sit between the cuticle ridges. Beyond 81% of development this remaining actin network shows evidence of dissociation (S8C,F), beginning at the peripheral margins of the cell. This suggests the actin network may be withdrawing from the cell upon completion of cuticle deposition. At 87.5% and beyond TauSTED imaging was not possible due to the presence of pigments.

The remodeling of the actin cytoskeleton throughout the later developmental stages follows the trajectory of cuticle deposition from the ridges to the crossribs (Fig 5). We also observed potential direct templating roles of actin in distinct ultrastructures including ridge layers, microribs and crossribs. At 63% development we observed a scale finger which was angled in such a way that a side profile of the ridge was visible. Directly below the cuticle ridge layers we noted layers of actin filaments (Fig 5Ai) which apparently matched the layering of the cuticle. At 69% we similarly observed a ridge side-profile which showed many small filaments of actin angled from the vertical along the side of the ridge (Fig B, Bi). This patterning of actin strongly resembles the final positioning of microribs along the ridges (Fig 1F). Finally, at 75% of development enlarged Z-stacks indicate that square blocks of actin form around the interior of the crossribs, though they do not fill the entirety of the nascent windows (Fig 5C, Ci).

Utilizing super-resolution microscopy during butterfly scale development, we have revealed new insights into actin cytoskeleton remodeling in butterfly scales. We have shown that the actin cytoskeleton plays a multifaceted role in butterfly scale development, from specifying finger location to a role in the development of ultrastructures, such as the crossribs and windows.

### The actin cytoskeleton plays a direct role in optical nanostructure formation

To determine whether actin plays a direct role in optical nanostructure formation, we injected pupae with Cytochalasin D (cyto-D), which inhibits actin polymerization and causes actin bundle disruption (*37*). Pupae were injected at 50% development, after ridge spacing is set but before ridge ultrastructures form and during incipient chitin deposition, to assess the effects of actin disruption specifically on structural color production (*21*).

We observed substantial loss of structural color in cyto-D treated forewings, with wings appearing visibly darker in color (Fig 6A,B) compared to the non-injected left forewing of treated individuals (Fig 6A), and the right forewing of controls injected with Grace’s Insect Medium (Fig 6C,D). Reflectance spectroscopy confirmed a significant reduction in brightness (t-test, t=4.34, d.f.=33, p<0.001) and flattening of the peak reflectance curve compared to control wings (Fig 6E). From the individual spectra plots (S9), most cyto-D treated individuals exhibited a completely flat reflectance spectrum with no change in angular intensity (i.e., no iridescence) and so total structural color loss (S9).

**Fig 6:**
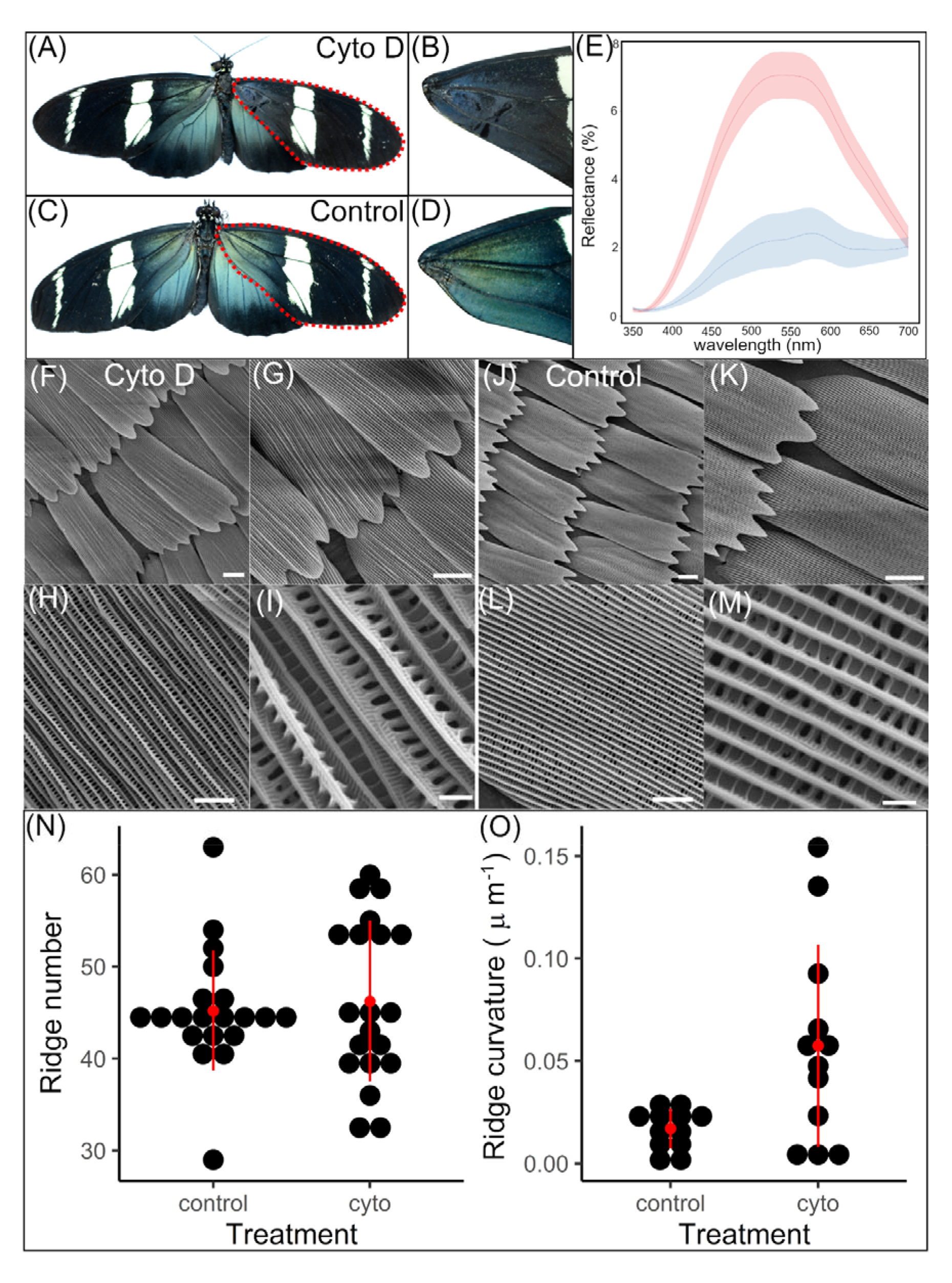
Chemical perturbation of actin with cytochalasin D at 50% development. Typical morphological phenotype of butterflies injected with cytochalasin D (A) and medium/DMSO (control, C) into the right forewing at 50% development and zooms of each (A→B, C→D) showing the discernible color change of the iridescent region. E) Reflectance spectra at the angle of maximum reflectance for control (red) and cyto-D treated (blue) wings. Shaded areas indicate standard error of the mean (for x measured individuals). F-M) SEM imaging of typical cyto-d treated (F-I) and control (J-K) individual’s wing scales in the treated region at different magnifications. Differences in brightness of the ridges indicates differences in height. N) Ridge curvature (κ) for cyto-d treated and control scales. O) Ridge number for cyto-d treated and control scales. Black points indicate individual scales. Red points and lines indicate the mean and standard deviation respectively. Scale bars lengths: (F, G, J, K) 15μm, (H, L) 5μm, (I, M) 1μm.

We observed no discernible differences in the size of cyto-D treated scales compared to control scales (Fig 6F-G, J-K). In some extreme cases we observed deformation of scale shape, with flexing of the fingers outwards and a ‘pinching’ of the central ridges (S10). There was no difference in the average ridge number between cyto-D treated and control scales (t-test, t=-0.41, df=5, p=0.70, Fig 6O). This confirms that by 50% development, ridge number and position has already been established in scale cells.

SEM imaging of cyto-D treated scales revealed significant deformation of ridge structure compared to controls (Fig 6I,M). This includes loss in ridge uniformity, evidenced by severe curving and collapse of the ridges (Fig 6H,I and S10). In terms of alteration of the ridge layering, pivotal for controlling the reflected structural color wavelength and intensity, this is clearly seen in figure S10 (B,C). With the ridge lamellae having morphology like that seen in a typical non-iridescent scale (see Parnell *et al* (*29*) for more examples of *Heliconius* ridge structures). We noted an increase in ‘breakpoints’ apparent in ridge layers of cyto-D scales (S11A), again more characteristic of non-iridescent scales, compared to the more continuous ridge layering seen in controls (S11B). We also observed that in some cyto-D treated individuals, the window regions were entirely filled with cuticle (S10 B,E). To quantify ridge disruption, we compared curvature (κ) of the ridges between treated and control scales (Fig 6N). Cyto-D treated scales had significantly greater average ridge curvature (κ) (μm^-1^) compared to controls (mean ± SE curvature (κ), treated 0.0566±0.0018μm^-1^, control 0.0158±0.0006μm^-1^; t-test, t=-2.78, df =12, p<0.05; Fig 6N). We also noted a large distribution in the average curvature values of treated scales, consistent with the differing levels of scale disruption observed in SEM images.

These results show that perturbation of the actin cytoskeleton during ridge formation results in significant loss of structural color. This can be directly attributed to the disruption of the scale ultrastructural elements responsible for iridescent color production.

## Discussion

The gross adult morphology of iridescent and non-iridescent *H. sara* scales does not differ dramatically, showing that only small changes are needed to produce structural color. However, our results show that iridescent scales have a substantial decrease in the spacing of parallel ridges. Through comparisons between developing iridescent and non-iridescent scales of *H. sara,* we determined that the reduced ridge spacing associated with adult iridescent scales can be attributed to a denser packing of actin bundles during development. Although a relationship between actin bundle spacing and ridge spacing has been shown previously (*20, 21*), we show that this association holds for structural color producing ridges. A tighter ridge spacing is crucial for maximizing reflectance and therefore iridescent scale properties (*35, 38*). As the layered lamellae responsible for iridescence in *H. sara* are present within these ridges, closer ridge spacing increases the density of light-reflecting surfaces within an individual scale. In the butterfly *Morpho adonis*, (which also contains layered lamellae optical nanostructures), a reduction in ridge spacing of just 0.13 μm yields a 30% increase in reflectivity (*38*). Our results show that the actin cytoskeleton is crucial for controlling close spacing of ridges in iridescent scales, through denser packing of actin bundles during the scale development.

The developmental control of total actin bundle number within scale cells warrants further investigation. *Drosophila* bristle studies have highlighted several actin-binding proteins that may be key regulators of actin bundle abundance (*23, 39*). Perturbation of two such proteins, actin-binding protein 1 (Abp1) and Scar, within developing *Drosophila* bristles resulted in extra bristle ridges. These may be promising candidates for future studies of butterfly scale formation (*40*).

Dinwiddie et al., (*21*) observed that structurally colored, silver scales of *Vanessa cardui* possessed double bundles of actin between ridges. In contrast, we observed very little difference in bundle organization between iridescent and non-iridescent scales of *H. sara* (Fig 2). This is likely due to differences in morphology linked to structural color production. The iridescent scales of *A. vanillae* have fused windows, with chitin between their ridges, reduced crossribs, and highly patterned microribs. These significant differences in scale architecture are linked to the optical phenomena that *A. vanillae* harnesses to produce structural color; whose formation involves dramatic shifts in chitin deposition likely controlled by actin patterning (*21, 26*). In contrast, layered lamellae in iridescent *H. sara* scales are patterned onto an already existing structure – the parallel ridges. There is no dramatic shift in architecture between iridescent and non-iridescent scales in *H. sara* and therefore the actin organization is similar.

If the hypothesis proposed by Ghiradella (*16*) is correct and F-actin provides the stress forces necessary to induce elastic buckling of the cuticle layer into layered lamellae, then perhaps we should expect to observe differences in actin dynamics, such as compressive forces, rather than large-scale differences in organization. Indeed, our perturbation of the actin cytoskeleton using Cytochalasin D and the resultant dramatic reduction in iridescence (Fig 6A) support a more direct role of F-actin in controlling the layered lamellae architecture. Cytochalasin D promotes sub-bundling of actin, resulting in wavy and distorted actin bundles within cells (*41, 42*). We saw that disruption of actin bundles and therefore mechanical integrity during optical nanostructure formation causes considerable reduction in iridescence (Fig 6).

The deformed ridges observed in our cytochalasin D treated butterfly scales (Fig 6I) display similarities to bristle phenotypes observed in fly mutants for actin organization proteins (*27, 43, 44*). As for fly bristles, actin bundles in butterfly scales are crucial for ridge formation, which occurs through extracellular chitin deposition in the inter bundle regions (*24, 27, 36*). Without actin bundles correctly guiding these projections, the final chitin ridges form in an aberrant manner, leading to ridges of varying geometries (*27*). We see loss of structural color in our treated samples in part attributed to collapse of ridges into varying angles, resulting in the multilayer photonic nanostructures no longer in registry with one another and therefore preventing concerted light reflection.

Interestingly, we observed additional phenotypic effects of actin perturbation on ridge ultrastructure. Harnessing both SEM (S10) and AFM (S11) we noted regular ‘breakpoints’ appearing on the usually continuous ridge layers. In these images we also see that the lamellae in the ridges have a strong variation in their thickness, this again points to the underlying reason for the loss in photonic properties (Fig 6I,M). The disruption of ridge layering suggests a further role of actin in directly controlling the formation of layered lamellae. Whether this perturbation of actin disrupts the stress forces needed to buckle the cuticle into layers, as predicted by Ghiradella (*16*), or instead prevents correct localization of chitin synthase enzymes to deposit the ridges (*26*) presents an interesting topic for future investigation.

Cytochalasin D may also have disrupted the secondary branched actin network present within scales (Fig 4I,L). Our TauSTED imaging showed that this network was particularly prominent after 63.5% development, when the cuticle ridges had already formed, and the parallel actin bundles were breaking down (Fig 4G-I). We speculate that this network may be involved in stabilizing the scale cuticular structures as the prominent parallel actin bundles break down. During this stage, the scale is still filled with cytoplasm and therefore likely subject to high cytoplasmic pressure (*45*). In support of this prediction, we see actin filaments in between the cuticle ridges as well as a high density of branched actin at the scale edges and in the fingers (Fig 4,5). Furthermore, some scales treated with cyto-D exhibited loss of overall uniformity, such as splayed fingers, consistent with disruption to a scale-wide stabilizing mechanism (S10A). The branched actin filaments may act as a series of intracellular ‘struts’, keeping the complex cuticular ultrastructure in a fixed registry until cuticle deposition is completed. Interestingly, at 75% development the actin becomes ‘block-like’ as it arranges around the crossribs (Fig 5C), suggesting that this stabilizing mechanism of actin may follow the path of the depositing cuticle internally as scale development progresses.

In conclusion, our study shows that the actin cytoskeleton plays a crucial role in the development of structural color in the butterfly *H. sara*, and likely the many other species that produce color through similar ridge reflector structures. Through denser packing of actin bundles during development, iridescent *H. sara* scales attain a higher density of chitin ridges enhancing the optical reflectance. In addition, using actin perturbation experiments, we demonstrate that the actin cytoskeleton likely plays a more direct role in the development of layered lamellae. The actin “scaffold” appears to template the chitin deposition and make the chitin structures (as they are forming) more mechanically stable during this process. Absence or diminution of the actin results in photonic structures that are out of registry with one another and also disruption of the lamellae that comprise the Bragg reflective layer, leading to substantial changes in the overall reflected intensity and directionality of the structural color.

We postulate that the role of actin may be akin to the layout and pinning stage used in dressmaking, so it is crucial to achieving high levels of long-range order and perfection across an entire scale cell. Ultimately, a better understanding of how the actin cytoskeleton controls structural color development in butterflies will help us understand how such complex natural photonic structures evolved and are patterned within individual cells. This has broader implications for our understanding of intracellular patterning more generally and for the design of synthetic systems to produce materials with similar optical properties.

## Materials and Methods

*Butterfly rearing -* Stocks of *Heliconius sara* were established from pupae originally purchased from Stratford-upon-Avon Butterfly Farm, United Kingdom. Adult butterflies were maintained in breeding cages at 25 □, and fed on 10 % sugar water solution with ∼1 gram of added pollen per 200 ml. *Passiflora auriculata* was provided for adults to lay eggs on. Caterpillars were kept at 25 □, 75 % humidity and fed on *Passiflora biflora* shoots. Pre-pupation caterpillars were checked regularly, and the time of pupation was recorded as the point of pupal case formation.

Wing scale development occurs during the pupal stage (*46*). At the desired stage, wings were dissected from pupae in phosphate buffered saline (PBS) and immediately fixed for 15 minutes in 4 % paraformaldehyde in PBS, at room temperature. Developmental stages of pupae were recorded as a percentage of pupal development, with *H. sara* taking 8 days from pupal case formation to eclosion at 25 □.

### Electron Microscopy

Adult wing samples were cut from regions of interest and adhesive tape was lightly applied to remove some cover scales. Samples were sputter coated with gold before being imaged on a JEOL JSM-6010LA SEM, equipped with InTouchScope software. See SI for TEM methods

### Immunofluorescent microscopy

Fixed wings were stained with various combinations of: mouse Anti-□-Tubulin primary antibody followed by a Cy3 AffiniPure Donkey Anti-Mouse secondary antibody for microtubules; Phalloidin or SiR-actin for actin; Wheat Germ Agglutin (WGA) for membrane and chitin, which was later replaced by Chitin Binding Domain that is specific for chitin. Slides were stored at 4°C until imaged. For each slide, both the proximal iridescent region and distal non-iridescent region were imaged. Confocal microscopy imaging was performed on a Nikon A1 confocal laser microscope equipped with NIS elements software. Super resolution imaging was performed on a Leica TCS SP8 STED microscope with Falcon module (see SI for details).

### Comparative analyses of iridescent and non-iridescent scales

10 males and 10 females, were used for SEM analysis of adult scale morphology with 10 cover and 10 ground scales analyzed for each individual. 12 pupae at 50% development were used for phalloidin staining to measure actin bundle number, size and spacing, with 5 scales measured in each wing region (blue vs black). Image analysis was conducted in ImageJ (*47*) (See SI for details).

### Chemical perturbation of actin

Actin inhibition experiments followed the protocol of Dinwiddie et al., (*21*). Ready-made cytochalasin D solution (Merck) (5 mg/ml in DMSO) was diluted to a final concentration of 20 μm in Grace’s insect medium (Merck). Pupae were injected at 50% pupal development using a Hamilton microliter syringe (701N). 5 μl of drug was injected directly into the proximal portion of the right wing blade. Control pupae followed the same protocol but were injected with 5 μl of 20 μm DMSO in Grace’s insect medium. Pupae were allowed to continue development until eclosion. Immediately after the wings had dried post-eclosion, butterflies were humanely killed. Butterflies which failed to emerge properly were discarded from further analyses. Only batches with an eclosion rate of over 50% were included in further analyses. A chi-squared test was used to assess differences in emergence rate between control and treated pupae. Whole wing imaging was performed on a Nikon D7000 DSLR camera. Scale imaging was performed using SEM and AFM (see SI).

### Statistical analyses

All statistical analyses were performed in R (Version 3.5.2) (*48*). For SEM analyses of adult iridescent and non-iridescent *H. sara* scales, we constructed a linear mixed effect model for each response variable (scale area, scale length, scale width, ridge spacing, ridge width) using the lme4 package (*49*). Prior to fitting the mixed effect model for ridge width, we averaged individual ridge measurements per scale. For models of ridge spacing, scale area and ridge width we included ‘individual’ as an intercept only random effect and for the model of ridge spacing, we included an interaction term between scale type and region. For scale length and scale width we fitted a random slope mixed model, allowing a different response to wing region for each individual. We used likelihood ratio tests between models with the Chi-squared distribution to assess statistical significance of sequentially dropped terms. For pairwise comparisons, Tukey multiple comparison tests were performed using the emmeans package in R (*50*). For analyses of the ridge spacing between the proximal and distal scales of *H. e. demophoon,* we firstly averaged measurements for each region per individual. Given the lower sample size we performed a paired t-test.

For analyses of actin bundle width, bundle spacing and bundle number in developing iridescent and non-iridescent scales, we constructed linear mixed effects models using the Lme4 package. For bundle spacing and bundle width, we firstly averaged bundle measurements per scale to account for multiple bundle measurements. For all models we fitted ‘individual’ as an intercept only random effect and tested statistical significance using likelihood ratio tests with a Chi-squared distribution.

All figures were constructed with ggplot2 (*51*), GIMP (v.2.8.22.) (*52*) and ImageJ (*47*). See Supplementary Information for the R scripts and data used to undertake these analyses.

## Supporting information

Supplemental information; including methods and supporting data

Animated Z-stack of an actin bundles (green) in an iridescent scale at 44% development (Fig 4A-C). Dashed line indicates the outline of the scale cell

## Acknowledgements

We thank Dr. Alan Dunbar (CBE, University of Sheffield) for use of the scanning electron microscope (funded via the EPSRC grant EP/K001329/1). We are grateful to Chris Hill (Electron Microscopy Facility, University of Sheffield) and Frane Babarović (School of Biosciences, University of Sheffield) for assistance in transmission electron microscopy. Confocal microscopy was conducted at the Wolfson Light Microscope Facility, University of Sheffield, and we thank Darren Robinson for his assistance. We also thank Dr. Natalia Bulgakova and Dr. Miguel Ramirez (School of Biosciences, University of Sheffield) for provision of an Anti-□-tubulin antibody and their expertise in immunofluorescent staining. We thank Esther Garcia Gonzalez (Central Laser Facility, Oxford) for assistance in the processing of the TauSTED data. We thank William Hentley (School of Biosciences, University of Sheffield) for assistance in the high-magnification imaging of the wings. Finally, we thank Gonzalo Castiella Ona who provided invaluable assistance in the laboratory during his Erasmus funded internship.

## Funding

This research was supported by a Natural Environment Research Council (NERC) Fellowship to N.J.N (NE/K008498/1) and doctoral training partnership (Adapting to the Challenges of a Changing Environment, NE/L002450/1) studentship to V.J.L and a Human Frontier Science Program Research Grant (RGP0034/2021). O.H. was funded by the Biotechnology and Biological Sciences Research Council (National Biofilms Innovation Center, PI A.J.P.). TauSTED imaging was supported by the Science and Technology Facilities Council at the Central Laser Facility (Experiment 20130025). For the purpose of open access, the author has applied a Creative Commons Attribution (CC BY) license to any Author Accepted Manuscript version arising.

## Author contributions

V.J.L., N.J.N., R.L.C. and A.J.P designed the project. V.J.L. and A.J.P. conducted most of the data collection, with the exception of the X-ray nanotomography (S.F., E.L. and I.G.) and its rendering which S.L.B undertook, and reflectance spectroscopy analysis which J.E.R performed. J.H. designed the STED experiment together with V.J.L., V.J.L and O.H, acquired, whilst V.J.L processed the STED data. All authors contributed to data interpretation. V.J.L wrote the manuscript, with all authors contributing to manuscript revision and editing.

## Competing interests

Authors declare that they have no competing interests.

## Notes

### Competing Interest Statement

The authors have declared no competing interest.

