## Supplemental information; including methods and supporting data for "The actin cytoskeleton plays multiple roles in structural color formation in butterfly wing scales"

**Supplementary information, figures, and tables**

***Supplementary results***

There was a significant difference in the length of iridescent and non-iridescent cover scales (Tukey comparison, p < 0.001), but not ground scales (Tukey comparison, p = 0.99) (mean ± SE scale length, blue (cover, ground) = 91.7 μm ± 0.48, 86.6 μm ± 0.47; black (cover, ground) = 97.4 ± 0.56 μm, 86.4μm ± 0.41). Correlation function analysis of the X-ray nano-tomography measured scales indicates a greater cross-rib spacing in the black scale compared to the iridescent scale (iridescent 0.483μm; non-iridescent 0.607μm)(S1). An expanded cross-rib spacing in black scales allows more light to enter the scale and so be absorbed by melanin pigments (*34*).

To confirm that differences in ridge spacing were not the result of sampling from two different wing regions (proximal vs distal) we measured ridge spacing in the closely-related, non-iridescent species *Heliconius erato demophoon* (S6). Ridge spacing did not significantly differ between the proximal (comparable to *H. sara* iridescent region) and distal (comparable to *H. sara* non-iridescent region) forewing (mean ± SE ridge spacing, proximal = 0.906 μm ± 0.0101, distal = 0.956 ± 0.0125 μm; paired t-test, t = 1.94, d.f. = 4, p = 0.125; S6 C-D). This shows that the observed differences in ridge spacing in *H. sara* are associated with the presence of iridescent structural color and not dependent on scale position on the wing.

***Supplementary Methods***

*Synchrotron X-ray nanotomography*

Individual scales were mounted vertically onto fine needle tips using a 3-axis optical alignment stage and UV-curable adhesive. X-ray nanotomography was performed at the imaging beamline at PETRAIII operated by the Helmholtz Zentrum hereon at the Deutsches Elektronen-Synchrotron facility (DESY), Hamburg (*53*). The samples were measured in a transmission X-ray microscope (TXM) at 11 keV in Zernike phase contrast mode, with an effective pixel size of 22.8 nm. Samples were rotated by 180° while acquiring projections. The tomographic reconstruction was performed in Python using Tomopy (*54*). A machine learning approach was used to denoise the 3D volumes after tomographic reconstruction (*55*). Axis alignment, thresholding and 3D volume reconstruction was performed manually in Python, using High Speed Tomography in Python (PyHST) software. See (*56*) for full rendering methods. Values of ridge spacing and cross-rib spacing were extracted from the reconstructed images using correlation function analysis.

*Immunofluorescence*

Fixed wings (4% PFA in PBS (phosphate buffered saline)) were washed several times in PBSTx (0.5-1% % Triton-X 100). For microtubule staining, samples were then blocked in 5 % Goat serum in PBSTx, rocking at room temperature for two hours. Primary antibody labeling was conducted using a mouse Anti-ɑ-Tubulin antibody (T6199, Sigma) at 1:1000 in PBSTx, overnight at 4 ℃. Secondary antibody incubation used Cy3 AffiniPure Donkey Anti-Mouse (Jackson ImmunoResearch) at 1:300 and samples were incubated at room temperature for 2-3 hours. For staining of the actin cytoskeleton / membrane / cuticle wings were left overnight at 4 ℃ in Phalloidin (Alexa Fluor 555/ ATTO-647; Invitrogen) or SiR-actin (Spirochrome) at 1:200 in PBS and/or Wheat Germ Agglutin (WGA) (Alexa Fluor 647 conjugate; Invitrogen) at 1:300 in PBS. For the STED microscopy dye concentrations were increased by 2-5 fold and WGA was replaced by a Chitin Binding Domain (TMR) at a 1:75 – 1:100 dilution in PBS (New England Biolabs; special request). Finally, DAPI (1 μg/mL) was used for counterstaining. Wings were mounted onto slides with Fluoroshield (Merck) for confocal microscopy and Mowiol or Prolong Diamond Antifade (Invitrogen) for STED microscopy and a coverslip applied. Left hindwings were used as controls, following the above protocol but omitting the fluorescent dye and/or primary antibody.

*Transmission electron microscopy (TEM)*

TEM preparation followed the protocol of (*57*). A section of the iridescent region from an adult *H. sara* wing was dissected and the ventral scales removed with sticky tape. Samples were washed for 30 minutes with 0.25M sodium hydroxide and 0.1% Tween. Samples were transferred to a 2:3 solution of formic acid / EtOH for 2.5 hours and dehydrated with 100% EtOH for 30 minutes. Samples were washed briefly with Propylene oxide and transferred to Epon epoxy resin with progressive washes of 15%, 50%, 70% and 100% Epon; with each wash lasting 24 hours. Then samples were embedded in resin moulds and cured in a 60 ◦C oven for 24 hours. A Leica ultramicrotome was used to cut thin sections of the sample (70-100nm) which were then stained using Uranyl acetone for 10 minutes before being washed twice with distilled water for 5 minutes. Further staining was performed using a solution of Lead nitrate, sodium citrate and 1M sodium hydroxide, followed by two washes in distilled water. Imaging was performed using a Phillips CM100 Transmission electron microscope at an accelerating voltage of 100 Kv.

*Atomic force microscopy (AFM)*

AFM imaging of wing scales was undertaken using a Digital Instrument Dimension 3100 Scanning probe microscope equipped with a Nanoscope IV controller. AFM was performed in tapping mode as previously described (*29*). Data visualization and image reconstruction was performed using the freely available software Gwyddion (*58*).

*Confocal microscopy*

Confocal microscopy imaging was performed on a Nikon A1 confocal laser microscope equipped with NIS elements software. Z-stacks were assembled into single images using FIJI (*47*).

*TauSTED microscopy*

Super resolution imaging was performed on a Leica TCS SP8 STED microscope with Falcon module. Post processing of images was performed in Huygens Professional. Images were firstly stabilized for lateral drift and the signal-to-noise ratio (SNR) was estimated for each image with acuity mode on. A conservative deconvolution strategy was selected for the estimation of initial values and the recommended CMLE (Classic Maximum Likelihood Estimation) was used as the deconvolution algorithm. Background was automatically estimated using a 0.7 µm area. A widefield search mode was used to identify the most in focus plane for background estimation. The PSF (point spread function) was estimated using a conservative optimization approach. The estimated parameters of background and PSF were then used for the final deconvolution, which was performed using an optimized iteration mode with a default quality of 0.01 for the maximum number of iterations.

*Comparative analyses of iridescent and non-iridescent scales*

*SEM Analysis of adult scale H. sara morphology*

*H. sara* butterflies were taken from breeding stocks maintained at the University of Sheffield. 20 individuals, consisting of 10 males and 10 females, were used for analysis. For each individual a forewing was removed and a 5 mm x 5 mm section of both the iridescent region and non-iridescent, black region was used for SEM. For each region, images were taken of 10 cover and 10 ground scales.

ImageJ (*47*) was used to measure the length of each scale, taken from the distal edge of the socket to the distal tip of the scale. Ridge spacing was calculated using PeakFinder Tool (*59*) (S4). At the midpoint of scale length, a 90 ° transect was taken across the scale to measure its width and the numbers of ridges were calculated using the PeakFinder Tool. The average ridge spacing was calculated as scale width divided by total number of ridges. Scale area was calculated using scale length multiplied by scale width, using the rectangular area as an approximation of true scale area. Ridge width was calculated using the Ridge Detection plugin (*60*) (S4). For each individual, 5 ground and 5 cover scales per region were used to calculate ridge width. For each scale, a 600 x 600 pixel section from the middle of the scale was selected for analysis. Contrast and brightness levels were adjusted manually to optimal levels using FIJI (*47*). The Ridge Detection parameters were set as follows: Sigma = 3.10, Lower Threshold = 0.51, Upper Threshold = 3.40, Minimum Line Length = 40.00, Maximum Line Length = 0.00.

*SEM Analysis of adult Heliconius erato demophoon scale morphology*

*H. erato demophoon* butterflies were taken from University of Sheffield collections (captive reared individuals), with 5 individuals used for the analysis. For each individual a 5 mm x 5 mm section from both the proximal forewing and distal hindwing were mounted for SEM. For each region images were taken of 5 ground and 5 cover scales. Ridge spacing measurements followed the methods described above.

*Confocal analysis of actin in developing scales*

The dissected forewings of 12 individuals were phalloidin stained following the protocol above. Stained wings were mounted on slides and imaged using a Nikon A1 confocal. For each slide, several Z-stack images were taken from both iridescent and non-iridescent regions using a x40 oil objective lens. From each wing region, we selected 5 scales for analysis which showed minimal disruption and the least overlap with other scales. Images were converted into 8-bit grayscale images in FIJI. A 100 x 100 pixel section of the scale was cropped out and stacked using the sum slices. Actin spacing was calculated as the mean distance between the centre of the actin bundles. The Ridge Detection plugin (*60*) in FIJI was used to calculate mean actin bundle width. The Ridge Detection parameters were set as follows: Sigma = 1.22, Lower Threshold = 4.76, Minimum Line Length = 10.00, Maximum Line Length = 0.00. The upper threshold was set between 12.58 and 21.93 to account for slight differences in brightness between experiments. At least 4 ridges per scale were used to calculate mean actin bundle width. In addition, the total number of actin bundles per scale was calculated from 20 black and 16 blue scales from across 11 individuals.

*Reflectance spectroscopy*

Reflectance spectrometry was undertaken using an Ocean Optics USB2000+ Spectrometer connected to a PX=2 pulsed xenon light source with a fiber-optic probe. Right forewings from 15 controls and 21 treated individuals were mounted onto a rotating optical stage and measurements taken following the methods of Parnell et al., (*29*). SpectraSuite (Ocean Optics) software was used to acquire scans. Integration times were set at 350 ms, using 5 scans to average and a boxcar width of 3 nm. Data was analyzed in R, using the package PAVO (v. 2.4.0) (*61*). Spectra were smoothed using the ‘Procspec’ function and peaks extracted using the ‘Peakshape’ function. Average spectra for control and treated individuals were plotted using the ‘Aggplot’ function.

*Cyto-d manipulation*

Injections of cyto-D or Grace's Insect Medium (GIM)/DMSO (control) were administered into the right forewing region within the pupal case. In total 76 pupae were injected (42 cyto-D and 34 control) across 6 batches. A 70 % emergence rate was recorded (29 cyto-D and 24 controls), with no significant difference in emergence rate between cyto-D and controls (χ^2^ (1) = 0.0211, p = 0.884). Only butterflies with minimally damaged forewings were selected for further analysis (21 cyto-D treated and 15 controls).

*Phenotypic analyses of actin perturbation*

SEM was used to acquire ridge number measurements for 4 control and 4 treated individuals following the methods outlined previously. Five scales per individual were selected from regions across the sample. Statistical comparisons of ridge number between control and treated samples were analyzed using a Welch two-sample t-test in R. High magnification SEM images of three scales per individual were selected for curvature analysis. The Fiji package Kappa (*62*) was used to acquire measurements of curvature (κ) for 10 ridges per scale. Using control points plotted along individual ridges, curves were inputted as open B-splines and fitted using a ‘Point Distance Minimisation’ algorithm. The parameters were set as follows: Data Threshold Radius = 15, Global Threshold Level = 0.05 and Local Error Threshold = 0.05. All other parameters were maintained at default levels. Average ridge curvature was calculated per scale and comparisons performed using a Welch two-sample t-test.

AFM was performed on scales from the right forewing of 2 control and 2 treated individuals, using the methods outlined above. For each scale four transverse cross-sections were taken across the image encompassing as many ridges as possible and the profiles plotted. Several replicates were taken per individual. From these profiles, ridge height measurements were calculated using the absolute minimum value of the data to the top of the ridge peak. Average height per ridge was plotted for the treatment and control scales.

***Supplementary figures***

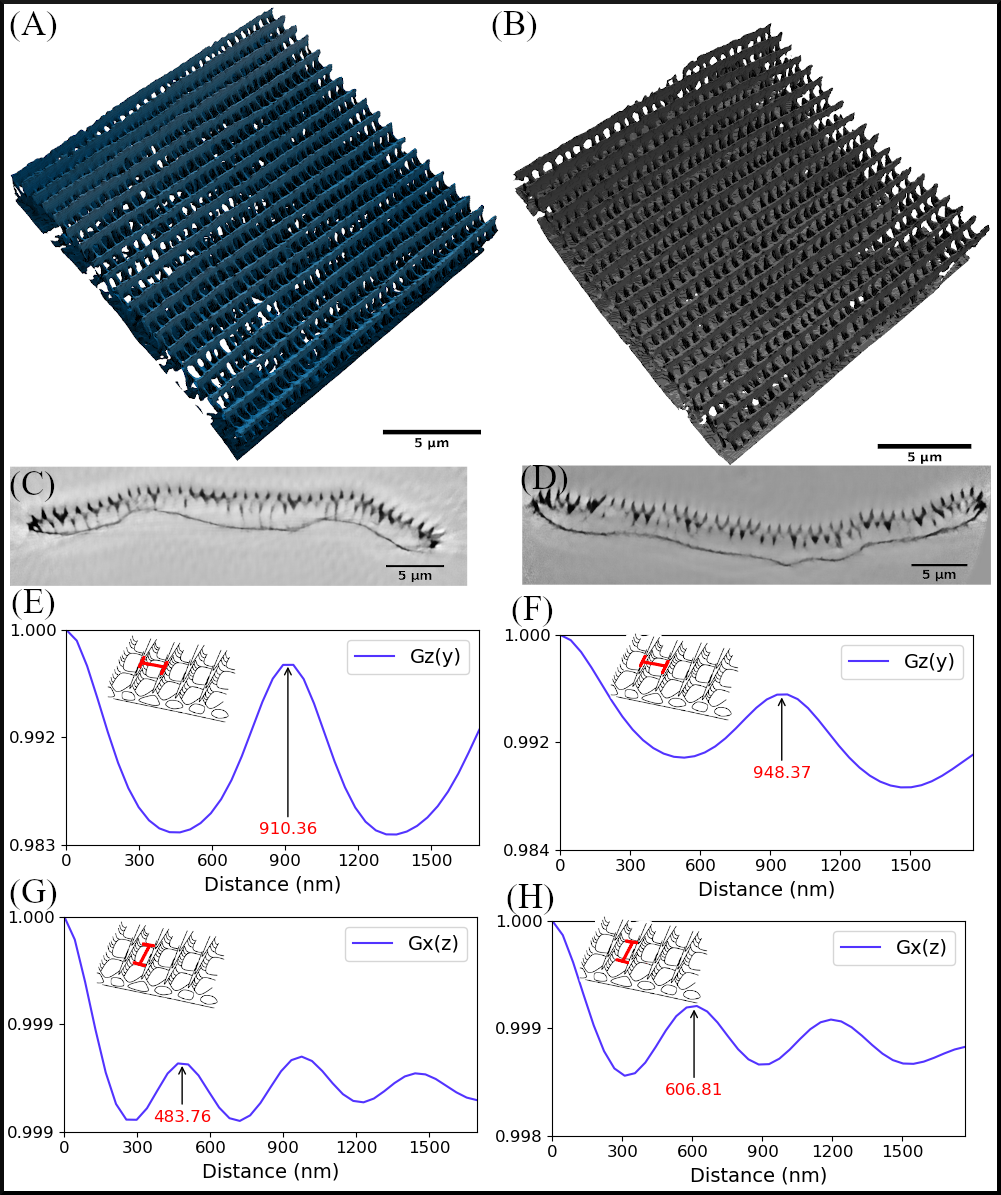
**S1: Synchrotron X-ray nanotomography of an adult iridescent and non-iridescent scale of *H. sara*.** Reconstructed 3D images of the central region of a blue, iridescent (A) and a black, non-iridescent scale (B) demonstrating the similarities in general morphology of both scale types. Sliced view (YZ plane) of the rendered iridescent (C) and non-iridescent (D) scales. (E-H) Extraction of morphological parameters from the rendered images using correlative function analysis. Red number indicates the measured parameter value. Extraction of ridge spacing from the iridescent (E) and non-iridescent scale (F). Extraction of cross-rib spacing of the iridescent (G) and non-iridescent scale (H). Scale bar lengths: (A, B) = 5 μm, (C, D) = 5 μm

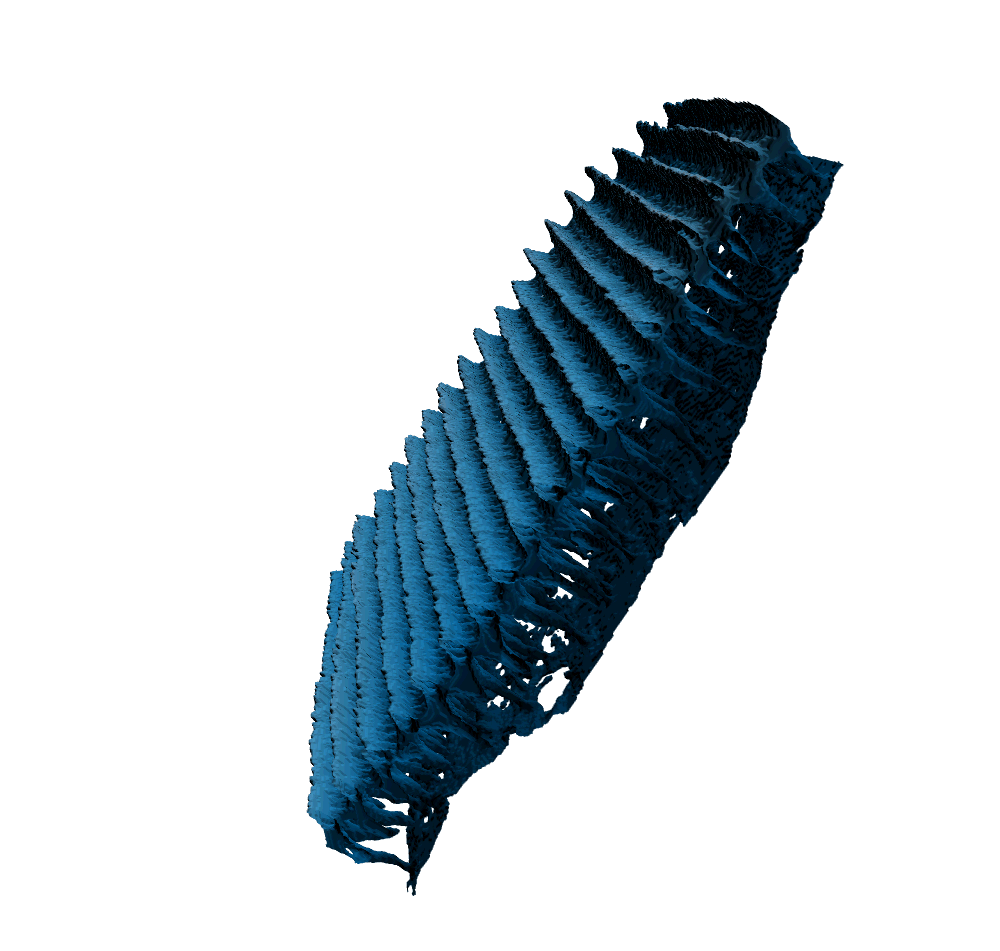

**S2: Movie of an X-ray nanotomography reconstructed blue, iridescent scale. False colored, blue.**

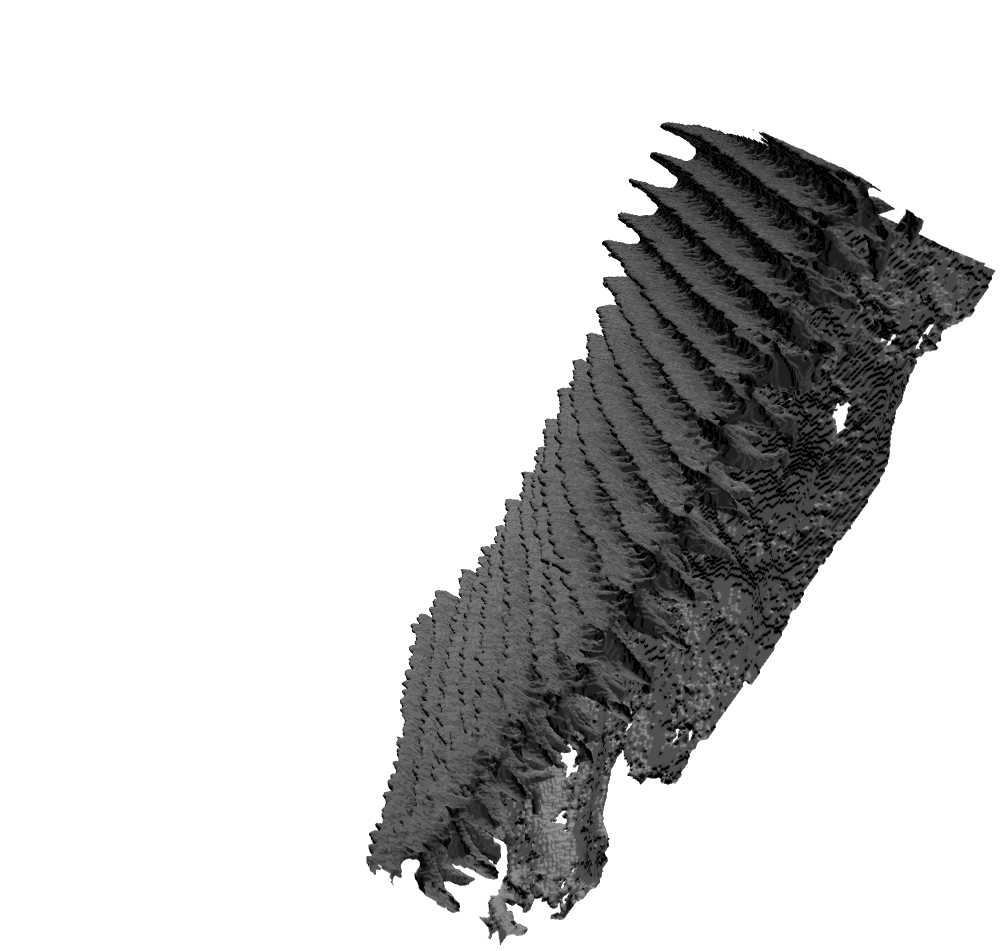

**S3: Movie of an X-ray nanotomography reconstructed non-iridescent, black scale. False colored, black.**

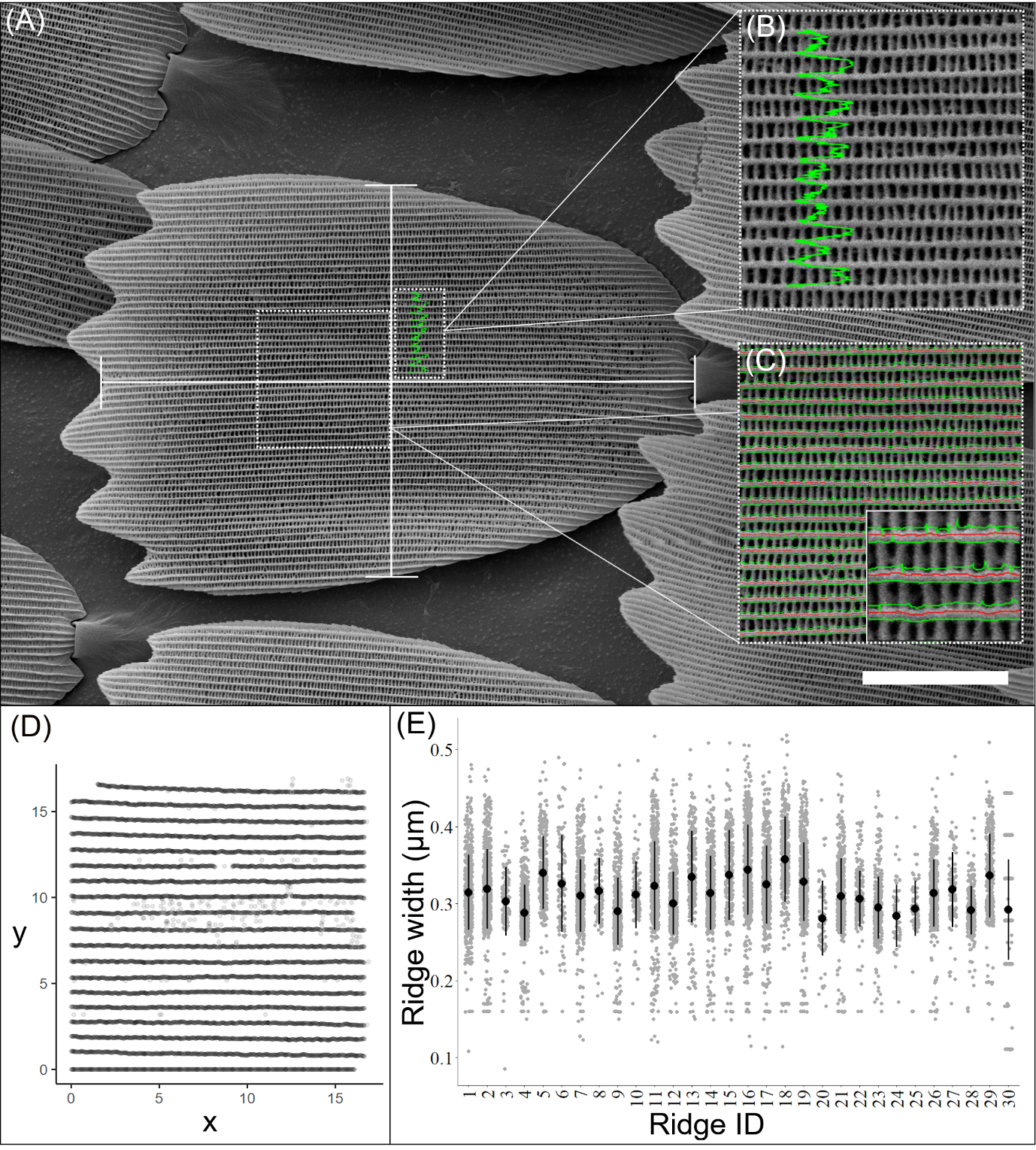

**S4: Methods of morphological analysis for an adult *Heliconius sara* wing scale.** A) SEM image of a *Heliconius sara* ground scale. The length of the scale is measured as the distance from the edge of the socket to the distal tip of the scale blade (horizontal line). The scale width is taken as a 90 ◦ transect at the midpoint of scale length (vertical line). B) PeakFinder tool can be used to select the longitudinal ridges from an SEM image. Each green peak highlights a separate longitudinal ridge, with the space between peaks forming a trough. C) The ridge detector plugin selects individual ridges (red lines) within the selected area of the scale. Each detected ridge is assigned a unique ridge ID. The green lines highlight the edges of an individual ridge and are used to calculate the ridge width. Measurements of ridge width were taken along the full length of each selected ridge. D) Positioning of all the 11,361 individual measurements of ridge width taken from the detected ridges in the selected scale area. E) All measurements of ridge width (μm), the mean ridge width (black dot) and standard deviation (black lines) for each selected ridge. Scale bar length = 20 μm.

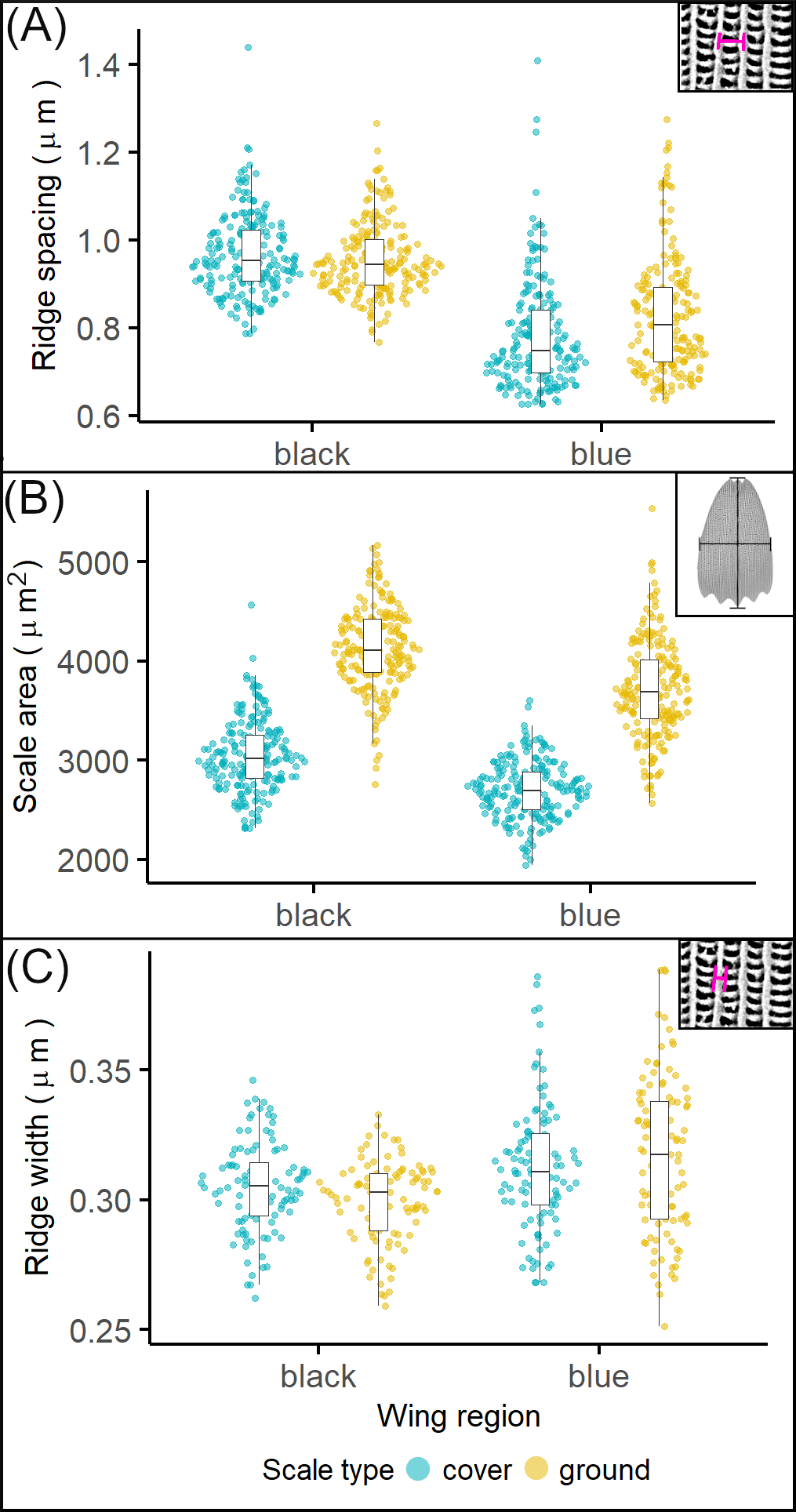

**S5: All data used for the SEM analysis of adult *Heliconius sara* wing scales.** Each data point represents an individual scale (n = 800). Measurements of ridge spacing (µm) (A), scale area (µm^2^) (B) and ridge width (µm) (C) taken from 10 cover scales and 10 ground scales in both the iridescent and non-iridescent region of 20 individuals.

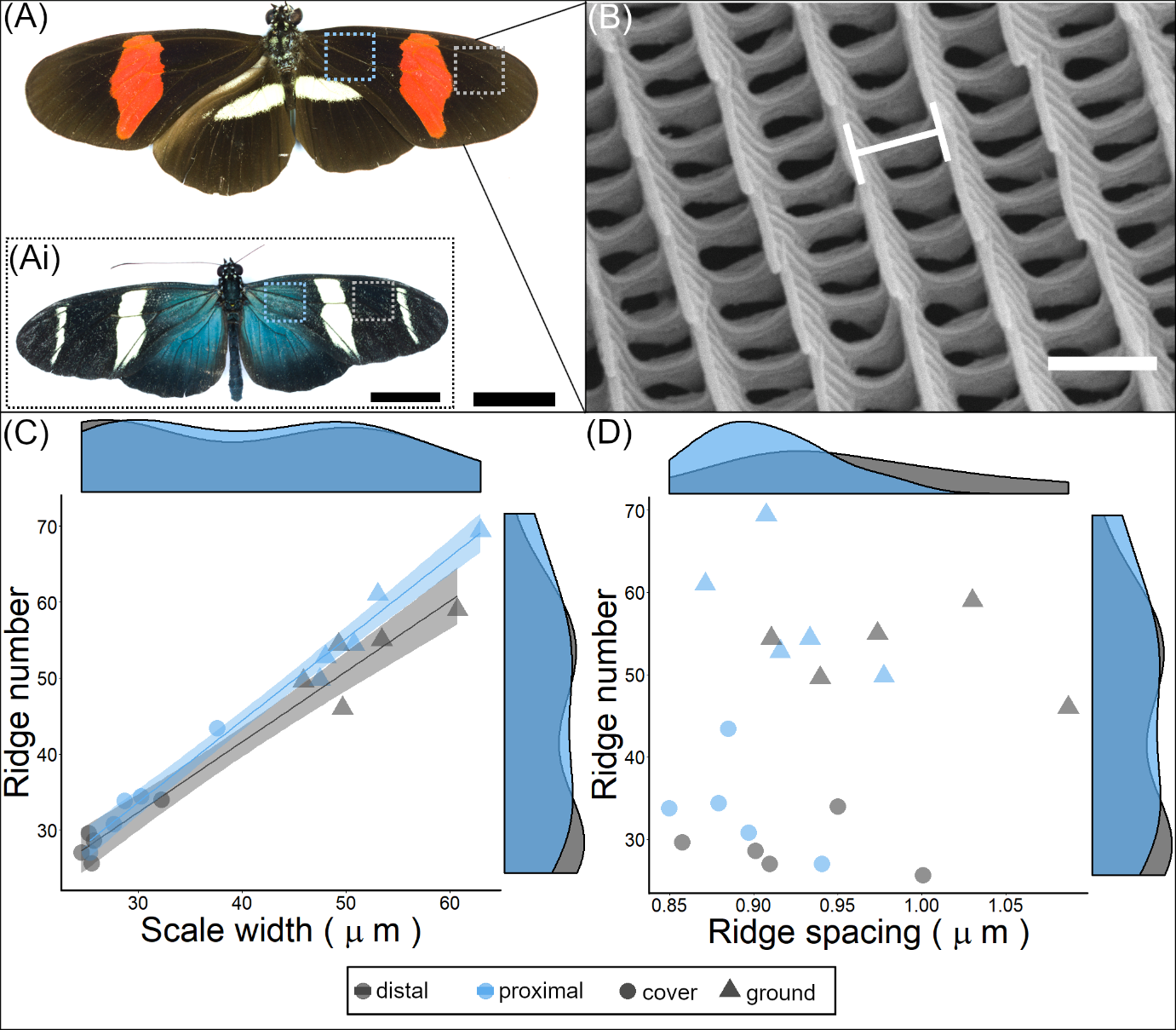

**S6: Morphological analyses of adult *Heliconius erato demophoon* wing scales.** A) *H. e. demophoon* individual with regions of interest taken from the proximal (blue box) and distal (grey box) forewing. These regions correspond to the iridescent and non-iridescent regions of *H. sara* (Ai). B) SEM image of *H. e. demophoon* scales ridges, white line indicates the ridge spacing. Comparisons of cover and ground scales in proximal and distal wing regions for (C) ridge number and scale width (μm), (D) ridge number and ridge spacing (μm). Each point represents the mean value grouped by individual, region and scale type. Shaded areas around the regression line indicate 95 % confidence intervals. Scale bar lengths: (A, Ai) = 30 mm, (B) = 1 μm.

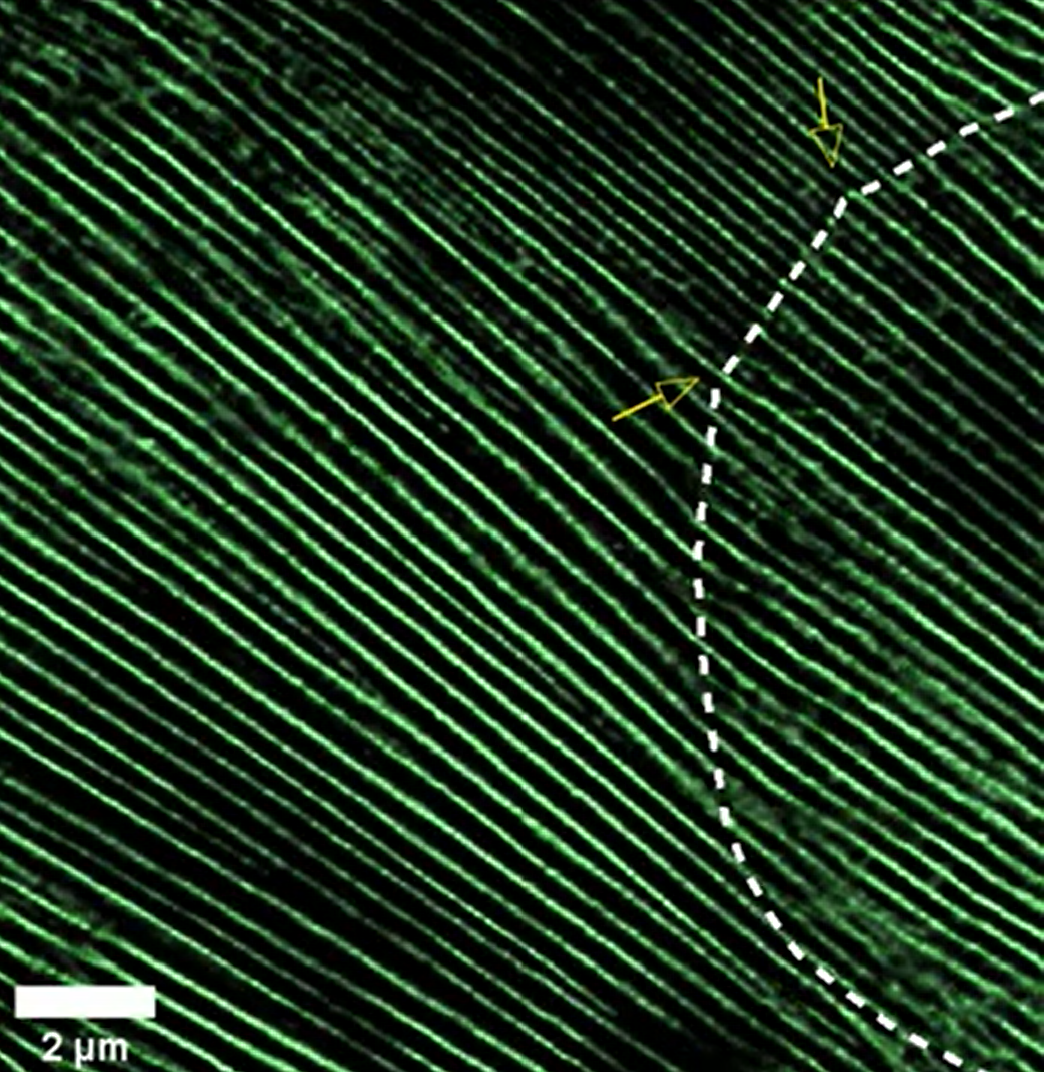

**S7:** **Animated Z-stack of an actin bundles (green) in an iridescent scale at 44% development (Fig 4A-C)**. Dashed line indicates the outline of the scale cell and yellow arrows correspond to the points at which finger formation begins on the distal cell edge.

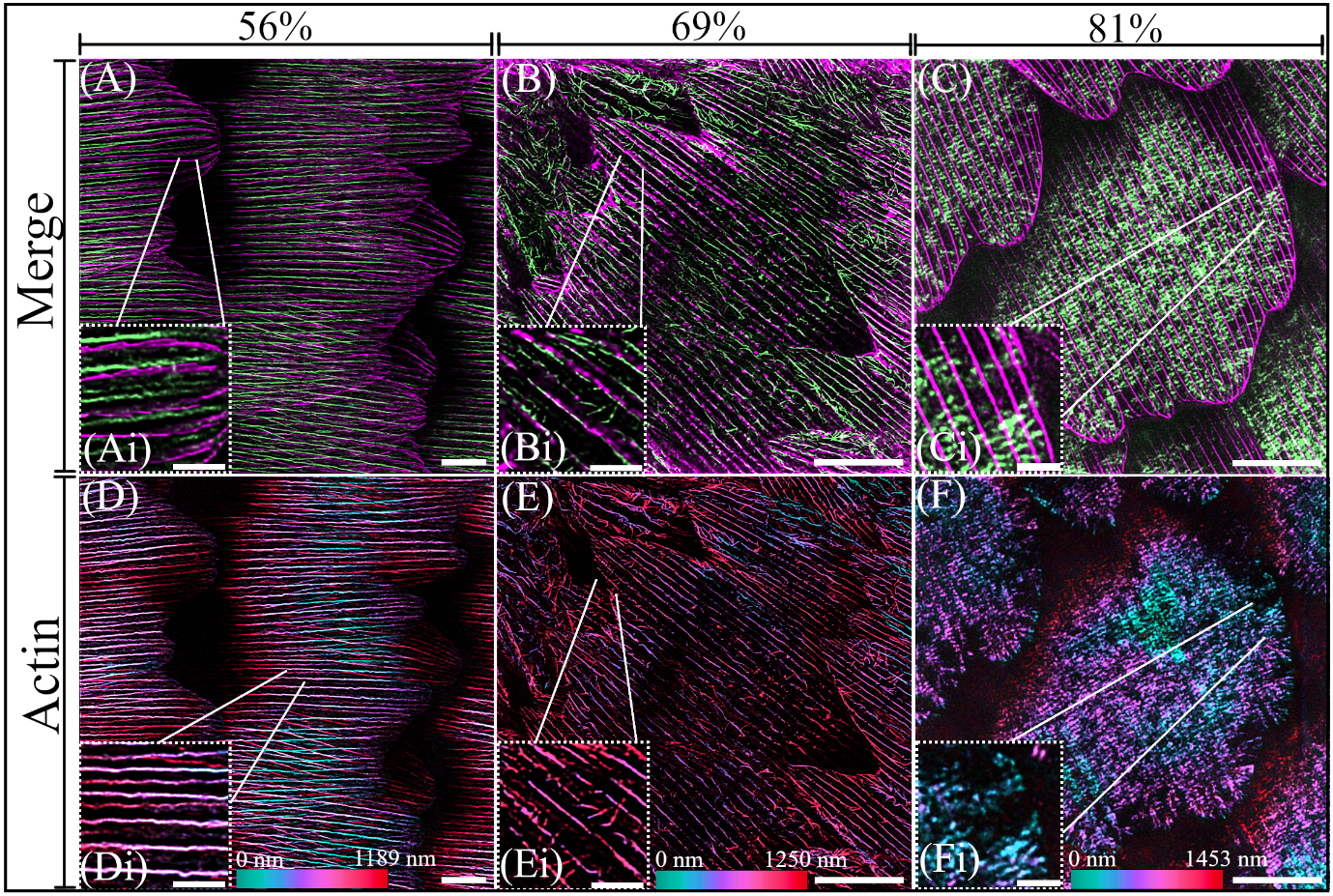

**S8: Additional TauSTED images of developing *Heliconius sara* scales.** (A-C) Merge of actin (green) and chitin (magenta). A) Scale from the black region of the posterior hindwing at 56%, with actin bundles positioned in between the chitin ridges. Individual overlapping layers of the cuticle ridges are clearly visible in the enlarged section (Ai). B) Iridescent scale at 69%, the large actin bundles are dissociating and actin filaments rearranging. (Bi) Enlarged section showing rearrangement of the actin bundles into individual filaments which now closely associate with the individual chitin ridges. C) Black scale from the posterior hindwing at 81%, showing the extensive deposition of cuticle and the final arrangement of the actin cytoskeleton before completion of scale development. In the enlarged section (Ci) the actin cytoskeleton can be seen withdrawing from the periphery of the cell. D-F) Depth color coded images of the actin cytoskeleton in images (A-C) with the enlarged sections showing regions approximately shown by the white lines. D) The actin cytoskeleton is present as large continuous bundles which appear relatively uniform in their Z position. E) The actin bundles fracturing and rearranging. Enlarged section (Ei) shows the rearranged individual filaments are more ventrally positioned than the original actin bundles. F) Final arrangement of the actin cytoskeleton. At this stage the actin cytoskeleton is less well defined at the periphery compared to 75% (Fig 5J) and regions lacking in actin can be observed. Together this suggests that the actin filaments may be beginning to break down entirely. (Fi) enlarged section indicates that the actin cytoskeleton is not present in large regions of the cell periphery. Scale bars: (A, D) 5 μm, (B, C, E, F) 10 μm, (Ai, Bi, Ci, Di, Ei, Fi) 2 μm.

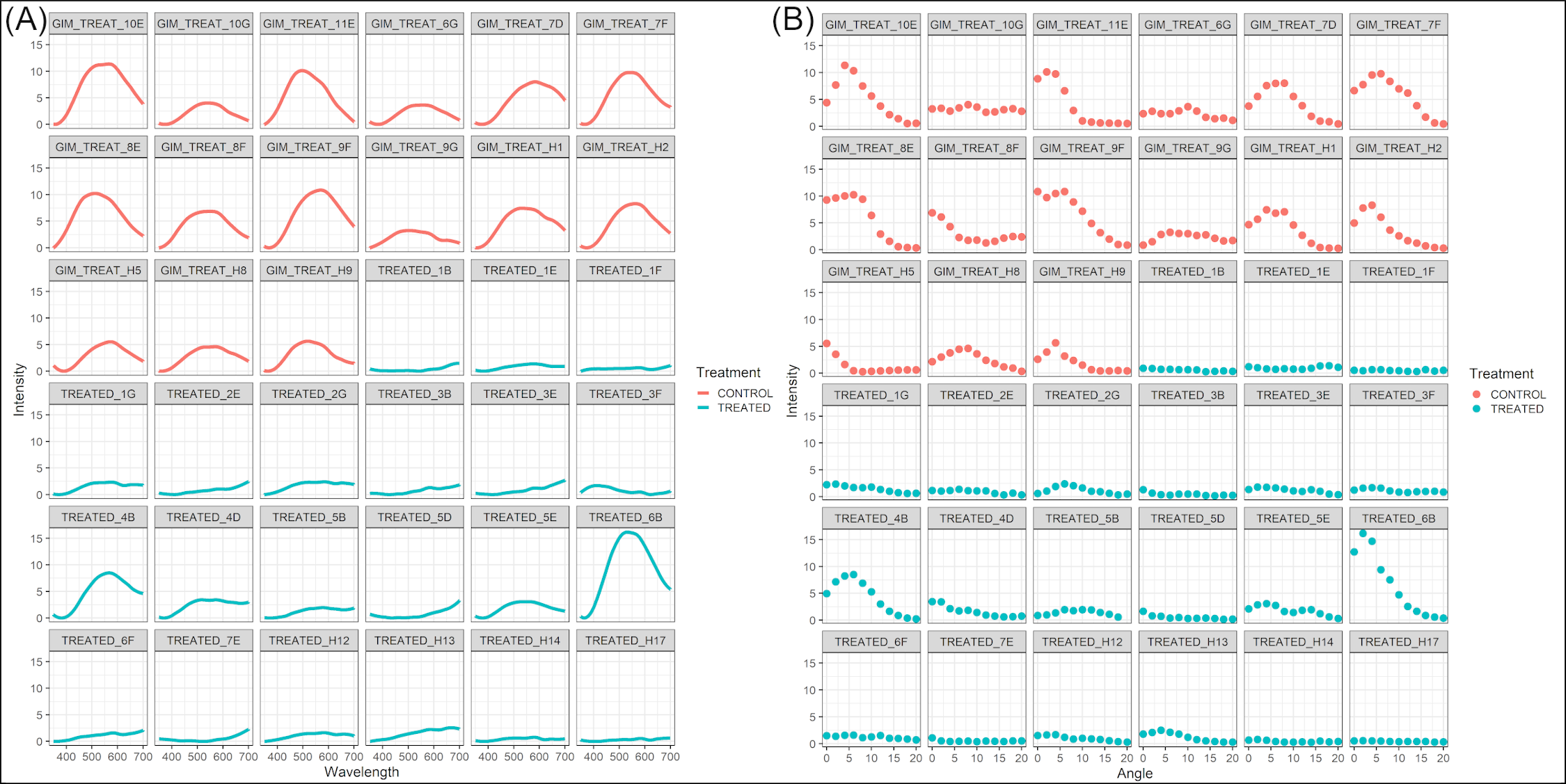

**S9: Individual reflectance spectra from all control and Cytochalasin D treated individuals**. A) Reflected intensity for measured wavelengths between 350 and 700 nm B) Reflected intensity for each angle measured between 0º and 20º.

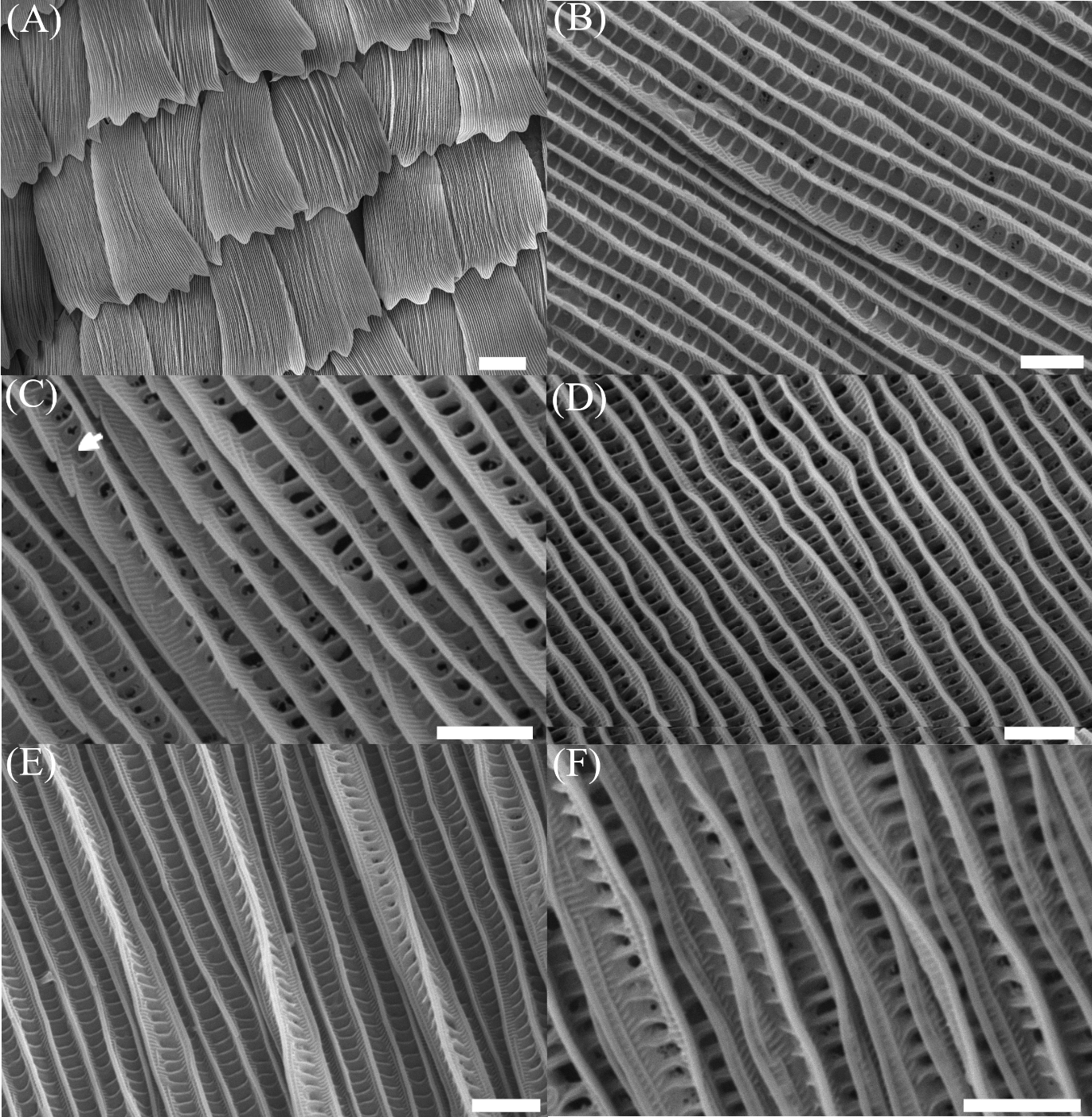

**S10: Additional SEM images of iridescent, blue scales from several butterflies treated with Cytochalasin D.** A) In some cases, defects of the whole scale were visible, with pinching of the ridges in the scale center, coupled with deformation of the fingers creating a fanned appearance of the scale tip. B) Some scales had windows entirely filled with chitin cuticle. C) Uneven layering of the ridges, with some ridge layers close together (white arrow) while other layers extended some distance. D) Acute curvature of some treated scales with a noticeable ‘wobbling’ of the ridges. E) In extreme cases, combinations of defects such as filled in windows as well as ridges toppling were seen. This was particularly the case in the ‘pinched region’ in the center of scales shown in (A). F) Some scales exhibited relatively normal ridge layering, but the ridges had toppled over and were laying out of plane with each other. Scale bars: (A) 20 μm, (B-F) 2 μm.

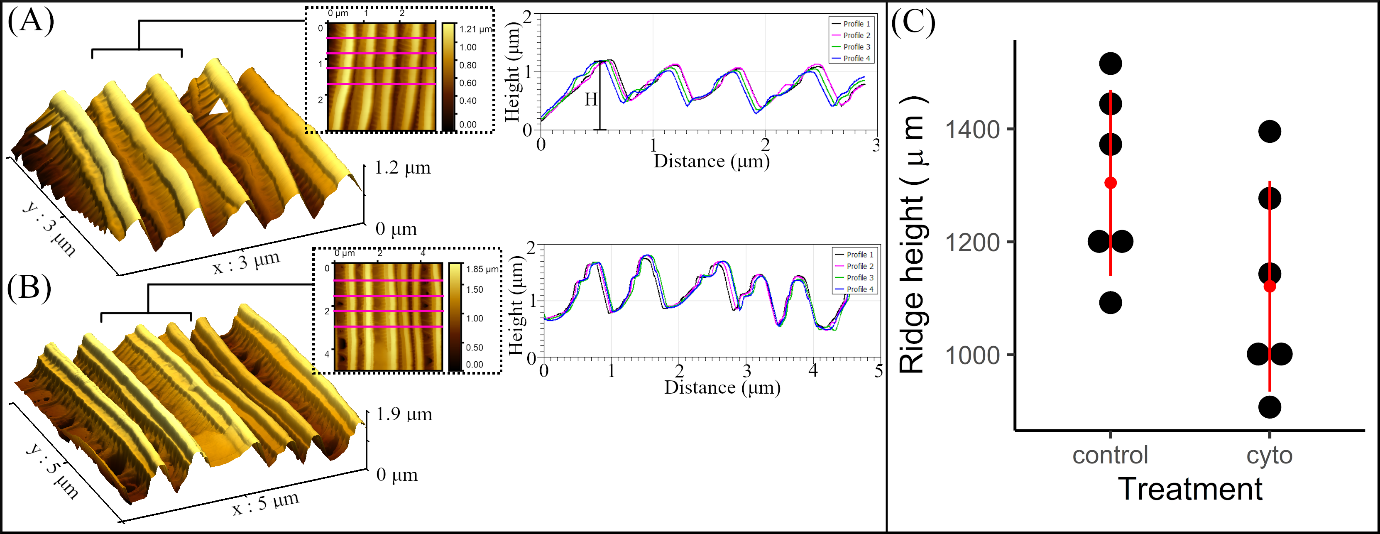

**S11: Atomic Force Microscopy (AFM) imaging and analysis of cytochalasin D treated and control scales.** 3D AFM rendering of scale ridges in a typical cytochalasin D treated individual (A) and a control individual injected with Grace's Insect Medium (B). Arrowheads in (A) indicate putative distortion of the ridge layering. Insert: AFM image of the 2D scale surface with magenta lines indicating method of selecting the four ridge profiles shown in the corresponding profile plot. Height (H) is taken as the distance from the peak of each ridge to the lowest point of the overall scale height. C) Plot of ridge height (µm) for control and cytochalasin D treated scales. Points represent individual scales.
